# Methods for assessing bee foraging preferences: a short review and a new automated apparatus

**DOI:** 10.1101/2023.12.15.571537

**Authors:** Evin T. Magner, Jeff T. Norris, Emilie C. Snell-Rood, Adrian D. Hegeman, Clay J. Carter

## Abstract

Bees are essential pollinators for many plant species, but multiple threats exist to both managed and wild bee populations. Their innate foraging behaviors and food preferences are subjects of intense research since bee nutrition is essential for maintaining hive and colony health. Multiple approaches have been developed to assess bee foraging behavior and associated preferences, but they are often labor-intensive and provide data on a limited number of parameters. In this manuscript, we provide a short review of these methods and present the design, build, and implementation of a new, inexpensive, and automated feeding apparatus capable of recording: (1) number of visits, (2) total food consumed, (3) photos and videos of bee behavior while inside of the feeder, and (4) environmental conditions. The efficacy of this apparatus is demonstrated through preference tests with artificial nectars, while also acknowledging the relative strengths and limitations of this approach.

## INTRODUCTION

Bees play essential roles in the pollination of plants in natural and agricultural systems(Dainese et al., 2019; Lemanski et al., 2022; Papa et al., 2022; Simpson et al., 2022). To improve the production of these systems and safeguard the survival of many flowering plants, we must understand the foraging habits and preferences of bees(Dainese et al., 2019; Garibaldi et al., 2013; Tong et al., 2023). Floral characteristics, including abundance, size, shape, color, patterns, volatiles, and nectar and pollen quality, influence the foraging preferences of bees (Balfour & Ratnieks, 2023; Lehrer et al., 1995; Nicolson, 2022; Su et al., 2022). Environmental conditions, hive/colony health, and nutritional needs can also affect bee preferences. However, due to their size, extensive ranges, and environmental factors, it can be challenging to study bee preferences for different food sources in wild and domesticated populations(Agatz et al., 2023; Campbell et al., 2019; Liira & Jürjendal, 2023; Mayack et al., 2023; Wagner, 2019).

Accurately counting bee visits and obtaining reliable quantitative data associated with their preferences has posed a challenge to researchers(Prendergast et al., 2020). Traditional sampling methods, such as manual counting, have proven time-consuming, labor-intensive, and prone to human error(Agatz et al., 2023; Prendergast et al., 2020). Prendergast et al. (2020) recently reviewed sampling techniques used for native bees, and Odemer (2021) reviewed automated bee counting systems. Below, we further examine a range of approaches, address the obstacles and limitations encountered, and point to recent advances that have contributed to the modernization of bee monitoring technology(Odemer, 2022). Most methods (**Table 1**) lack the ability to provide data on multiple parameters simultaneously, such as visitation rates, behavior and duration of visits, rate of consumption, and environmental conditions. We subsequently present a new apparatus that combines proven methods with newer technologies to study bee foraging preferences. The system incorporates technologies, such as Raspberry Pi computing, cameras, IR sensors, weight scales, and environmental sensors, to provide a comprehensive and automated data collection and analysis approach.

### A short review of methods to study bee preference

Various methods are used to investigate different aspects of bee foraging behavior and preferences, each posing distinctive strengths and limitations (**Table 1**). Visual observations, for instance, involve monitoring bees in a real-world environment, which provides significant insights into foraging behaviors(Leach et al., 2023). However, this approach has inherent limitations, including its vulnerability to subjective interpretation by observers and the inherent challenge of accurately tracking movement, decision-making, and species identification. Conversely, sweep netting involves systematically passing a net through an environment to capture bees, facilitating more accurate species identification. While this method can capture a wide variety of bee species, it may be limited by its reliance on chance encounters and often misses the foraging decisions of the bees.

Mobile gardens enable researchers to evaluate bee preferences in a range of environments and multiple locations by relocating plant material to different study sites for analysis(Lowenstein et al., 2015). It is important to note, though, that these gardens may not accurately reproduce natural environments and, therefore, potentially produce biased results. In natural habitats, the use of beating sheets as a method for detecting the presence of species within a specific geographic region, environment, or plant species may yield more accurate results(Avisar et al., 2023). However, depending on the research objective, this method may not adequately reflect the complexities of bee dietary preferences. In native environments, researchers also employ mark-recapture methods for studies with the ability to resample a population. Mark-recapture studies involve the practice of marking and monitoring individual bees, which may provide valuable insights into bee movements and preferences but incurs a high cost in terms of the resource-intensive nature and potential limitations in accounting for variation amongst populations(Harmon-Threatt & Anderson, 2023).

Passive collection techniques, such as the use of Malaise traps, sticky traps, soapy pan traps, and windowpane traps, have demonstrated efficacy in catching a diverse range of airborne insects, including bees(ATAKAN & PEHLİVAN, 2015; Skvarla et al., 2020; Visscher & Seeley, 2023). Still, these techniques are limited in their ability to offer comprehensive details regarding the preferences and foraging behaviors of bees. Preferences can be indirectly assessed using certain baits or targets. For instance, pan and bowl traps are designed to attract and capture bees by relying on baits or targets, rendering them well-suited for studying bee preferences based on color or other stimuli(McCravy & Ruholl, 2017). Although, the selection of an unnatural target might constrain the ability of the pan trap to measure natural foraging situations accurately.

In certain instances, bee preference studies may use more sophisticated technical approaches. Feeding platforms equipped with cameras deliver glimpses into bee behavior during feeding(Murphree, 2022). It is important to note that these open-air platforms may unintentionally introduce artificial factors, which could potentially bias the observations. Furthermore, although comprehensive, video recordings provide behavioral data that may necessitate substantial manual time and effort for subsequent data processing and analysis, thus rendering them less ideal for large-scale investigations(Gernat et al., 2023). For smaller-scale investigations, infrared (IR) sensors provide precise bee passage and visitation rate data, but their cost and complexity can pose challenges that may limit their widespread use(Pešović et al., 2017). In contrast, large-scale approaches use acoustic monitoring and Doppler radar, which excel at discerning bee flight patterns and activity levels but may not elucidate specific floral preferences (Sharif et al., 2022; Williams et al., 2023). Additionally, the implementation of Radio Frequency Identification (RFID) tracking technology developed for monitoring bees may have some limitations in revealing detailed feeding preferences while primarily focusing on following their movement patterns(Alburaki et al., 2021).

Alternative preference studies employ molecular and laboratory-based approaches to study bee preferences and foraging behaviors. One method, pollen analysis, highlights the types of pollen collected by bees, offering an opportunity to explore foraging choices(Layek et al., 2022). However, its focus on dietary aspects may not fully encompass behavioral preferences since it primarily determines what bees eat, overlooking other facets of preference, like color, shape, and scent. In contrast, controlled choice experiments conducted within the confines of a laboratory provide a systematic means to assess bee preferences(Mustard et al., 2019) but since laboratory environments differ from those of natural conditions, this approach may alter the outcomes and applicability of the findings. Electrophysiological recordings approach the issue from a different perspective by delving into neural responses to stimuli, providing valuable information about bee sensory perception(Ma et al., 2021). Though, establishing direct correlations between neuroactivity and behavioral preferences can prove challenging. Similarly, Y-maze or T-maze tests, designed for controlled choice studies, may not capture the entire spectrum of natural tendencies, potentially leaving certain subtleties unexplored(Nouvian & Galizia, 2019). Focusing more on the signal instead of its perception, chemical analysis of floral compounds offers invaluable insights into bee preferences, albeit with an emphasis on odor-based cues, potentially neglecting other sensory modalities such as visual or tactile stimuli(Fernandes et al., 2023). Another more straightforward approach with similarities is the Restricted Volume Preference Assay (RVPA)(Richman et al., 2021). This experimental procedure employs a compact box containing limited quantities of liquid or gas, from which the bees can make selections. Also, within the broad molecular-based toolset, Proboscis Extension Reflex (PER) experiments investigate bee responses to specific chemical stimuli but risk ignoring other elements of choice(Alqarni et al., 2023). Conversely, Free Moving Proboscis Extension Response (FMPER) assays offer a more natural testing environment in which bees have mobility, though they grapple with technical complexities and may not perfectly replicate outdoor conditions(Muth et al., 2018).

Genetic and molecular techniques likewise delve into the genetic underpinnings of bee preferences, shedding light on the role of genetics and evolution in bee foraging preferences. Yet geneticists may unintentionally overlook other influential factors guiding choices, such as colony-level behavior, competition, or even variation(Lüthi et al., 2022). An even broader approach is through phylogenetic analysis. Phylogenetic research investigates preferences among bee species and offers potential trends or correlations amongst clades, but the broad nature of the approach may not capture changes at the individual or population level(Vaudo et al., 2020). A computational alternative is through the use of mathematical modeling. Using a predictive approach, mathematical modeling depends on robust data for accuracy, making its outcomes contingent on data quality from the other monitoring methods described above(Liao et al., 2020).

Ultimately, each method described above has inherent advantages, disadvantages, and limitations (summarized in **Table 1**). Their appropriate usage is largely context- and question-dependent, such as: what is preferred? why is it preferred? how does preference vary? why does preference vary? and how is it perceived? Regardless of the context and questions being asked, we noted a general lack of methods that combine multiple sensors for the automated detection of bee visitation, consumption, behavior, and the associated environmental conditions. Our goal in this study was to develop an inexpensive modular system with the ability to integrate automated detection and measurement for each of these parameters.

## MATERIALS & METHODS – A NARRATIVE OF THE DEVELOPMENT OF A NEW APPARATUS FOR BEE PREFERENCE ASSAYS

### Overview

In order to overcome some of the constraints associated with the methods described above, we developed a new apparatus to assess bee feeding preferences by integrating Raspberry Pi computers with proven techniques, including extensive modification of dry pollen feeding tubes made out of polyvinyl chloride (PVC) tubes and fittings often employed by beekeepers(Anderson, 2023). Our newly developed Capture, Count, and Consumption (CCC) apparatus houses a feeder location, computers, and sensors inside of a similar PVC pipe (**Fig. 1**) and is connected to an external power supply located a short distance away, rendering the system portable and well suited for remote field studies. The system integrates a broad range of technologies, such as Raspberry Pi Zero computers, weight scales, infrared (IR) beam counters, IR motion sensors, and environmental sensors. By integrating these components, researchers can obtain reliable data, especially for social bee (e.g., *Apis* spp.) foraging preferences and visitation counts. This new apparatus can measure bee visitation rates while collecting additional parameters, including the duration of visits, rate of food consumption, video and images of bees, and environmental data. Integrating multiple technologies in one system enables a thorough comprehension of bee foraging and preference behaviors. Details of the design, parts, construction, and use are outlined below.

**Figure 1.**
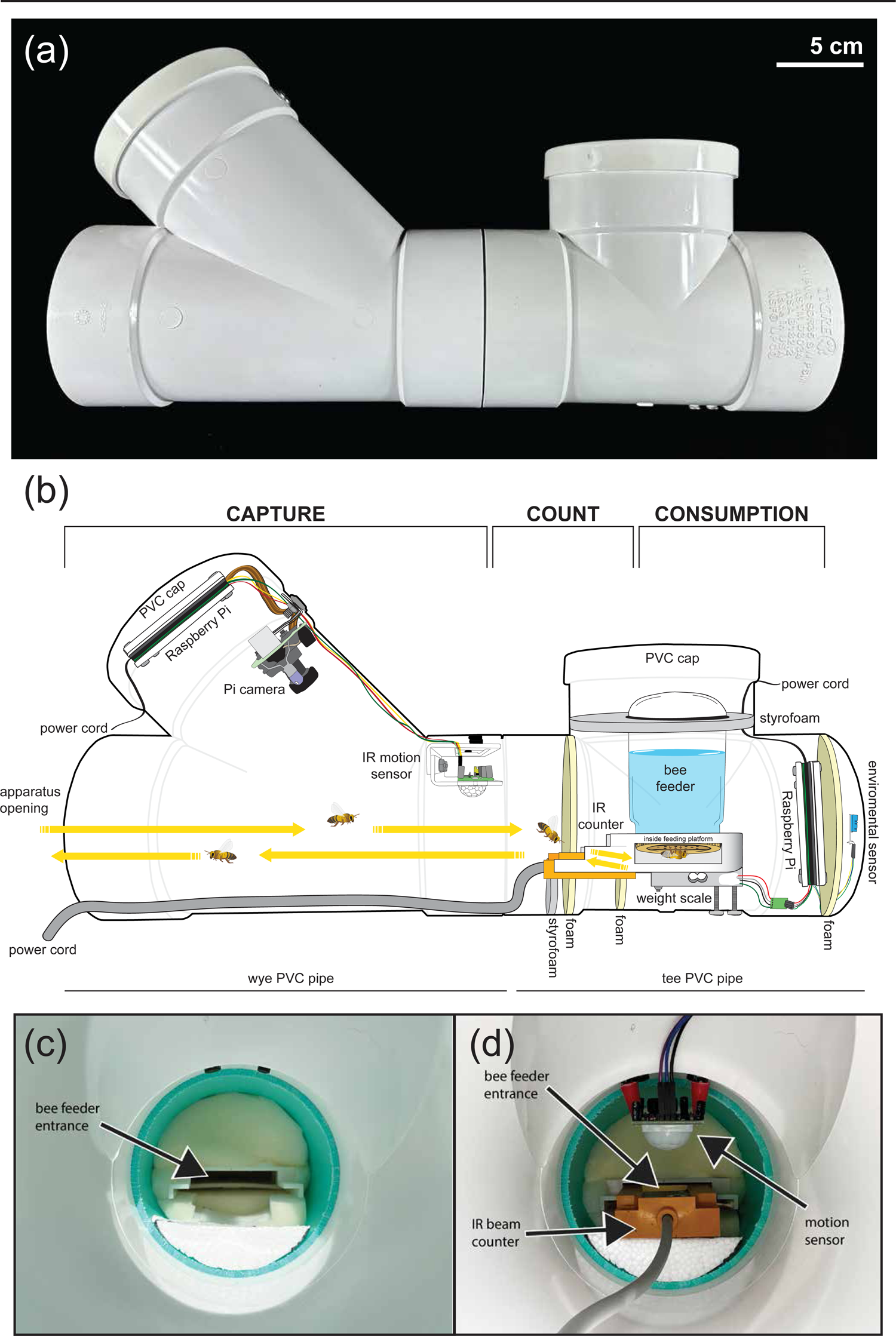
Design of the Counting, Capture, and Consumption (CCC) Apparatus. (a) Exterior view of the CCC Apparatus. (b) Longitudinal cross-section of the CCC Apparatus, showcasing the placement of various technologies and components within the tube. The labeled components provide a comprehensive outline of the design. (c) View from the entrance of the apparatus towards the back, revealing the feeding platform entrance. Components have been removed to enhance visibility, allowing a unobstructed observation of the opening leading to the feeding platform. (d) Similar to (c), but with the inclusion of additional features - an infrared (IR) counting beam and an IR motion sensor.

### Description of CCC parts and construction Apparatus housing

The primary housing unit of the CCC apparatus (**Fig. 1**) is based on that of simple pollen tube feeders employed by beekeepers (Anderson, 2023). In our design, a short section of 4-inch (∼10 cm) diameter PVC pipe (**part A.2 in Fig. S1A**) connects a tee (**A.3**) to a wye (**A.1**). A solid end cap (**A.4**) and a drainage grate (**A.5**) are attached to the top (short end) and sides of the tee section, respectively. A second solid end cap (**A.4**) is attached to the top of the wye, and an optional downspout adapter (not shown) can be added to the lower (straight) section of the wye to reduce the opening size of the apparatus. This lower section of the wye serves as the entrance point for bees, whereas the tee section is where the feeding platform is located (**Fig. 1B-D**), which also provides enough space for a Raspberry Pi (**B.1**), weight scale (**D.2**), and environmental sensor (**D.7**). A 0.5 pint (∼237 mL) mason jar attached to a nectar feeder with a perforated lid and an IR beam counter assembly (**C.1**) readily fits within the tee (**A.3**). The upper portion of the wye section houses a Raspberry Pi (**B.1**), a Pi camera (**B.2**), and an IR sensor (**B.3**), which activates the camera and can be used for direct counting. All electronics (described in greater detail below) are shielded inside the unit, ensuring their safety from the environment. All parts incorporated into the system are listed in **Table S2** and noted with their associated identifier from **Fig. S1** in the parenthesis following the part.

The housing unit offers a single entry and exit point. Bees access the apparatus at the lower section of the wye pipe and proceed through to the tee pipe, which houses the feeding platform and food source (e.g., nectar in this study) (**Fig. 1B**). Within the housing unit, bees trigger the camera by passing in front of a passive infrared (PIR) motion sensor (described in detail below). Once activated by the PIR, the camera has the capability to capture either video footage or still photos, depending on the specific needs of the research. Bees enter the feeding platform in the tee section of the unit and pass through the IR beam counter, which the digital readout records. The bees inside the feeding platform can consume the offered food, which the weight scale mounted to the feeding platform also records. During each weight assessment, the environmental sensors log the conditions during each weight measurement. Upon completion of feeding or investigating, the bees must return through the entrance of both the feeding platform and the housing unit, once again triggering the beam counter first, followed by the PIR motion sensor and camera. It is possible for more than one bee to enter or exit the feeder simultaneously, but we have no evidence that this characteristic alters the absolute counts or other measurements.

### Raspberry Pi configuration and code

This project uses multiple programs coded in the Python Integrated Development Environment (IDE) included on the Raspberry Pis (**B.1**) by default when using the Raspbian operating system. The complete code can be found in the specified GitHub repository [https://github.com/Jeff-Norris/Nectary_Project]. Software programs are required to operate the Pi camera, each of the Pi accessories, and potentially others to process results received from the former programs. If the intent is to replicate the same system as described in this manuscript, the code supporting the sensors should be utilized without modifications, assuming that the sensors are attached to the identical General Purpose Input/Output (GPIO) pins as those employed in this project (**Fig. S1)**. It is essential to activate and assign the correct pins in the code to avoid mistakes in transmitting and receiving messages between the Pi and connected devices.

Researchers can access the repository to examine, reproduce, and modify the code to suit their requirements. Our focus on promoting cooperation, developing research methodology, and contributing to the collective knowledge of the scientific community is in line with our commitment to open science. The repository functions as a consolidated source, providing both the code and accompanying documentation and assistance to facilitate the smooth implementation of these crucial procedures in various research environments.

### Passive infrared (PIR) sensors attached to Pi computers

This design has multiple options for employing passive infrared (PIR) sensors (**B.3**). First, one can use the sensor as a simple counting mechanism. Utilizing the PIR sensor as a counting mechanism requires a simple loop with the sensor waiting for motion and incrementing a counter variable every time detection occurs. For this to work correctly, one must ensure that the same entity is not causing the sensor to activate multiple times. This necessitates proper organization of the CCC Apparatus, as well as fine-tuning of the potentiometers of the PIR sensors (**Fig. S1B**).

The second option for using the PIR sensor involves activating the camera to begin recording a video or capturing an image that can be assessed later for bee counts or observation. Augmenting the primary loop from the option mentioned above can accomplish this. Instead of incrementing a counter when the sensor is triggered, the camera will capture until the sensor no longer detects motion, which will cease operation. By appropriately setting the potentiometers of the PIR sensor, one can reduce the running time, power consumption, and required storage for the video or images. However, there is a potential trade-off since discontinuous recording may result in missing information.

### Pi Camera

The Pi camera system located in the upper wye section of the apparatus (**Fig. S1B**) uses a Longruner Raspberry Pi camera. Further modifications added two Kingbright WP7113SF4BT-P22 infrared LEDs and aluminum heatsinks (primarily useful if recording visitation by nocturnal visitors in other studies). The *Camera.py* file in the repository is a basic illustration of how to develop a program to operate the Pi camera (**B.2**) for capturing both images and videos. Some modifications may be made to adapt the specific camera being utilized, as there may be slight variations among different camera models. Testing Pi cameras with varying sleep times is crucial, as some require more activation time. The sleep time in the code allows the camera a brief period to prepare for capturing pictures and videos. It is vital to consider this timing to avoid information loss or inefficient photo or video capturing.

The program also allowed us to specify either the number of pictures desired or the period between each, using the take picture function or the video length if taking a video. By default, the program is written to run endlessly and save the video at intervals equivalent to the value of the video length variable, but it may also ascertain the number of videos. The timestamp is appended to both images and videos and then saved in the user-specified directory.

### Weight scale and environmental data collection

The repository also contains the codebase necessary for completing measurements pertaining to weighing the feeding solutions (consumption by weight in grams) and recording environmental data. The measure of consumption occurs by weighing the feeding platform using a modified kitchen scale (**D.2** in **Fig. S1**), while environmental data is collected with a sensor (**D.7**) positioned in the rear of the tee tube **(Fig. S1D)**. The developed code, tailored to this experiment, is stored in the repository with the rest of the codes. The code consists of scripts aimed at measuring the mass of the remaining sugar solution and environmental conditions at regular intervals. Capturing these data involves incorporating precise weighing mechanisms connected to the environmental sensor that records temperature and humidity.

In brief, a scale is attached to the base of the feeding platform with a plastic spacer separating the units to equally displace the weight across the scale **(Fig. S1D)**. The scale is then mounted using threaded bolts such that it floats above the base of the tube. A supplied module connects the scale to the Raspberry Pi. At the start of each trial, the initial mass is recorded, and the subsequent consumption measured by comparing the mass differences. In concert with measuring the mass, environmental data is collected using an environmental sensor that logs the temperature and humidity. In our trials, this experiment collected environmental and mass measurements simultaneously every 30 minutes. However, this time between readings can be reduced significantly to capture more detail or can be linked to the PIR sensor and measured when activated. The versatility also extends to the placement of the environmental sensor. In our trials, we placed the environmental sensor in the rear of the device against the grated drainage cap. Conversely, the environmental sensor could be placed inside the housing to measure the ambient conditions inside the apparatus. In all, the placement of the sensors and utilized code can be readily modified to suit the needs and conditions of the varying investigations.

### GPIO pins

GPIO pins are crucial elements of the Raspberry Pi Zero (**Fig. S1**). These pins enable users to connect a wide range of valuable tools, such as IR sensors, weight sensors, HATs (hardware attached on top), and pHATs (PiZero HATS). Due to the varying roles of its 40 pins, it is crucial to have a clear understanding of the connections and functions of these pins. Incorrect usage of the pins can lead to potential harm to both the attachment and the Pi. The pins can be configured for either input or output. Sensors or buttons provide digital signals to the inputs of microcontrollers, while L.E.D.s, relays, or other devices are regulated using the outputs. In order to provide power to the attachments, the Raspberry Pi contains several ground connections, as well as two pins each for 3.3 volts and 5 volts of direct current (D.C.) power. The Raspberry Pi Zeros typically have a complete 40-pin board that can be securely attached to the corresponding ports. The entire pin board is numbered 1 through 40 and arranged in two rows. The left row has pins with odd numbers, while the right row contains pins with even numbers. To connect the board to most accessories, one will need to use female-to-male cables, although depending on the source of the component, this requirement may vary **(Fig. S1)**.

Alternatively, one can directly solder male connectors into the respective pin ports. Our attention will be solely on the essential pins required for connecting the sensors in this project. However, interested parties can find complete material regarding the pins and their respective roles on <u>Raspberrypi.com</u>. We use the PIR sensor connection as a brief example of connecting sensors to the Raspberry Pi. The PIR sensors possess three connections: VCC, GND, and OUT. Either pin 2 or pin 4 should link the VCC, which serves as the power supply connection, and the 5V pins. The GND pin, representing the ground, should be linked to one of the five ground pins. Usually, pin 6 is the preferred choice. Finally, when the sensor senses motion, the OUT pin generates a low-voltage signal to the Raspberry Pi. This can be linked to any of the designated pins on the board that are not power or ground pins. We utilized GP1, specifically located at pin 12.

### Independent IR beam counter assembly at entrance of feeder platform

The IR beam counter (**C.1 in Fig. S1C**) is an adapted piece of equipment regularly used in industrial settings for counting parts as they pass on a conveyor belt, for example. In our system, the IR beam is positioned at the entrance to the feeding platform. The sensor bar fits inside the feeding platform such that bees must pass through the beam to enter the feeding chamber. In order to focus the path of the bee towards the entrance of the feeder, Styrofoam (E.4) and rubber foam (E.1, E.3) were used to obstruct the back of the apparatus and support the IR in the opening of the feeder, as depicted in **Fig. 1C, D and Fig. S1E**.

Unlike other sensors within the system, the independent IR beam counter acts as a standalone component not linked to the Raspberry Pis. Instead, the IR beam counter connects to its own dedicated power supply and digital readout. In much the same manner as the other components, the design of this portion of the apparatus attempted to protect the electronics while providing ease of use and data acquisition. To accomplish this, a plastic waterproof ammunition box (**C.5**) was used to house a 12 VDC battery supply (**C.3**), digital data readouts (**C.2**), and power switch (**C.4**). In short, a single ammunition box provided the location for the power supply and digital readouts for two independent IR sensors (i.e., it can be used in binary choice preference assays). In practice, this single unit was used to collect data from two CCC apparatuses located in close proximity to one another (described in greater detail below). Researchers can manually collect the counts displayed on the digital readout before resetting them.

### Feeding platform and reservoir

To investigate food and foraging preferences, a food source, such as a sugar solution (artificial nectar), can be used. The current apparatus has been designed to test various liquid solutions; however, the system has the capacity to use other food sources that also incorporate other pollinator cues (e.g., color, scent) by making some minor modifications to the modular system (e.g., some options are provided in **Fig. S2)**. In the design tested, we used a honey bee hive entrance feeder (**D.1**), typically used to supplement water during droughts. The design, though, works seamlessly with our system as it provides both a feeding platform and a reservoir tank. The feeding platform is designed to allow bees to enter the chamber and feed directly from the reservoir.

Conveniently, the reservoir is simply a 0.5 pint (∼237 mL) mason jar with a perforated lid that is inverted and placed on top of the platform (**Fig. S1D)**. To support the glass jar without inhibiting the measurement of the mass, a Styrofoam ring (**E.5**) was cut and placed in the top portion of the tee pipe. The ring maintained the jar in an upright position without interfering with the ability of the jar to be weighed, allowing for both ease of consumption and data collection. Finally, as mentioned previously, the feeding platform was mounted to the scale to allow the measurement of consumption by weight.

### Environmental sensor

Environmental factors play a pivotal role in influencing bee foraging preferences(CORBET et al., 1993; Descamps et al., 2018; Prasad & Hodge, 2013; Russell & McFrederick, 2021a; Sandoval-Molina et al., 2020a). Therefore, the CCC Apparatus integrates an environmental sensor that collects temperature and humidity data to investigate a correlation between the preferences and the local environmental conditions (**Table S2** and **D.7** in **Fig. S1**). To better understand the influence of environmental variables on bee foraging behavior, it is vital to analyze both environmental data and bee visitation data or consumption rates simultaneously. Simultaneously examining the environmental data alongside bee visitation or consumption rates enhances the understanding of the influence of environmental factors on foraging behaviors and preferences.

### Power supply

As mentioned previously, a dedicated 12 VDC battery powered the IR counter beam. A second remote 12 VDC battery powered the remaining electronics to ensure uninterrupted operation. The implemented battery provided sufficient power for extended periods, minimizing the need for frequent maintenance and recharging. The Raspberry Pis and connected sensors shared this common power source, which did not require significant power. Our experiments found that a Might Max Battery ML50-12 12V 50AH S.L.A. (not included in the list) provided enough power to operate 4 Raspberry Pi systems (including 2 CCC Apparatuses) and the associated sensors. The battery was contained within a waterproof plastic enclosure, positioned a short distance from the two CCC units for which it provided power. Power cords were run from the battery unit to each Raspberry Pi through the solid plastic caps positioned on the top of both the wye and tee portions of the PVC pipes (**Fig. S1)**. Depending on the settings used, it could also be conceived that these devices could potentially be powered by small solar panels depending on the remoteness of the study site, or smaller dedicated battery packs designed specifically to power Raspberry Pi units.

### Field setup and testing of the CCC apparatus

In order to assess both the functionality and limitations of the apparatus, we performed a pilot study on bee food preferences, which collected the following types of data: images and videos of visitors, the number of bee visits, consumption rates, and environmental conditions. In our pilot study, two CCC apparatuses containing different artificial nectars were placed at a distance of approximately 10 m from a cluster of *Apis mellifera* hives located on the Minnesota Agricultural Experiment Station of the St. Paul campus at the University of Minnesota (44.988213 N, −93.177695 W) (**Fig. S3**). U-bolts were used to mount two CCC apparatuses to a 2-inch x 4-inch framing lumber board attached to two studded T-posts at a height of approximately 1m (**Fig. S3A**). The openings of the CCC apparatuses were directed inward, facing each other, at a distance of approximately 0.5 m. Adding this spatial dimension to the experimental design contributed to the evaluation of the performance of the apparatus in a real-world context. The goal of the setup was to provide space for free flight while maintaining close proximity to the feeders to limit additional outside factors influencing choice. Agricultural and horticultural fields harboring a variety of plant species surrounded the field site, providing alternative foraging choices for the bees. The power supplies and the digital readout for the IR beam counter were positioned between the adjacent T-posts (i.e., on the ground in between the two apparatuses).

A sugar solution or tap water were offered as choices to bees for us to evaluate the usefulness of the system. As a control, one apparatus contained only 200 mL of ambient temperature tap water, while the other system contained 200 mL of a 20% (w/v) sugar solution comprised of 10% sucrose and 10% glucose in tap water. The solutions were prepared fresh daily before placing the apparatuses in the field. Both systems were positioned in the field each morning for four consecutive days at approximately the same time [09:00 CST] and remained in the field for 6 hours before retrieval [15:00 CST]; however, the positions of the feeders were switched halfway through the experiment at noon [12:00 CST]. At the conclusion of each day, all equipment was washed using a diluted soap mixture to remove any remaining residues or pheromones left in the system. This approach was taken to help ensure that the previous day’s visits caused no residual influence.

Before the start of this experiment, bees were initially lured and acclimated to the apparatuses using two yellow plant labels with lemon oil applied to them. Each label was mounted to the 2-in x 4-in piece of lumber near the opening of each apparatus. The labels were removed after a few days of exposure while we adjusted the technical aspects of the apparatus. The following week, formal data collection began. At the start of each day, a random number generator randomized the position of each apparatus to either the east or west position on the post. The assignment of either the sugar solution or water, as well as the specific IR beam counter, was likewise randomized to avoid any biases arising from technological factors or remaining sugars. At the 3-hour mark each day, the sugar solution and water position were switched to eliminate the positional effects, and current IR beam counts were collected before the digital readouts were reset.

### Machine learning

Images and videos were processed using Adobe Photoshop and Adobe Premiere Pro. The images and video were analyzed manually before exploring automated machine learning (ML) techniques. The analysis was to both explore the counts and investigate if other species visited the feeders. Having tried two techniques of ML processes using two different sets of tools, some are better for the simple purpose of counting as this is a relatively simple task and can be accomplished with more straightforward tools, which are typically preconfigured to an extent; whereas, tools for the verification of species can be much more difficult. For an attempt at the latter task, we used Darknet. Darknet is an open-source neural network framework created using C and Cuda. It is fast and computationally efficient, making it ideal for real-time object detection. Unfortunately, by default, the detector is good at recognizing basic objects but must be trained for specific circumstances if the accuracy of the recognition software is to be trusted. This training process can be very involved, and accompanying it with the installation and necessary configuration of the tools are a potentially significant hurdle for anyone looking to implement them. While Darknet is efficient, it is still a very computationally heavy program and requires the installation of both Cuda and Cudnn (if one is using Nvidia) in order to run at its peak performance. Installation of these tools and their required dependencies are documented on the Nvidia website. These two add-ons are for parallel processing and maximizing efficiency for neural networks, respectively. Upon installation of these three tools, one must create a training set of images that highlight things that can be expected in the video from the CCC, which can then be used to train the program before running it against either video or still images. The more images and the better their quality, the better the training of the data. In the coming months, we hope to have a cache of images to train the program for bee recognition, which will subsequently be posted to the aforementioned Github for use by the broader community.

Data collected by the IR beam counter and the Raspberry Pis and associated sensors were collected and analyzed using Prism 10. For visitation counts, the recorded values were divided by two to account for the in and out pass of each bee. The values were subsequently log_10_ transformed into a more accurate comparison, and a t-test compared the total number of visits over a four-day period to check for a significant difference. A Chi-squared test was employed to compare the count results from each trial and test for significance. The measure of consumption by weight was evaluated using a paired t-test to look for significant differences. Finally, we investigated the correlation between consumption (by weight) and temperature through Pearson and multiple variable correlation analyses.

## RESULTS & DISCUSSION

### Capture mechanisms evaluation

We began by evaluating the performance of the PIR capture mechanism for triggering counts, videos, and photos. Comparing the number of triggering events of the PIR (a proxy for total visitor numbers) to the images taken and the standalone IR beam counter numbers showed that while the PIR sensor was triggered by honey bees, using the sensor attached to the Pi solely as a counting mechanism was insufficient. The PIR sensor did not work well as a standalone counter due to an intrinsic lag time between the camera and sensor, in combination with the large internal volume of space allowing for multiple bees to be in the space simultaneously, the sensor was unable to provide accurate counts. However, the PIR sensor consistently detected bee movements by successfully triggering the camera to capture images and video. The reliable functionality of the PIR sensor coupled with the camera suggests its effectiveness in accurately identifying the presence of bees. In brief, the sensor demonstrated its ability to identify the presence of bees and trigger the camera system.

The camera performed exceptionally well in capturing clear images and videos of bees inside the tubes. The quality of the captured media (**Fig. 2B**) allowed for the analysis of bee behavior, contributing to the overall success of the apparatus in recording relevant data. The images also demonstrated that only honey bees visited the apparatuses during our trials. Images proved to be a more efficient use of storage space than videos and were adequate for analysis. However, videos provided a more detailed examination of bee behavior at the cost of storage space. In concert with the PIR sensor, the camera and trigger counts could illustrate the frequency of visits, even though the counts were inaccurate (see above). In order to overcome this deficiency, researchers can manually count bees in images or videos to obtain accurate counts. While time-consuming, manual analysis of images and videos may be the most accurate approach in ascertaining species-level identification and the duration of the visit. We also explored using ML to identify and count bees, producing mixed results. Simpler ML tools provided a clear picture of where bees were in still frames (**Fig. 2C**) but were not able to extrapolate this information in the next frame to differentiate a new bee from one which has already been counted, leading to inaccurate results. More robust ML tools seem to be able to counter act this issue with an added cost of training and customizing the tool itself. Ultimately, even with the challenges of incorporating ML and having multiple bees in the image space concurrently, the capture mechanism was shown to be functional and efficient and validated the system for monitoring bee activities in field settings.

**Figure 2.**
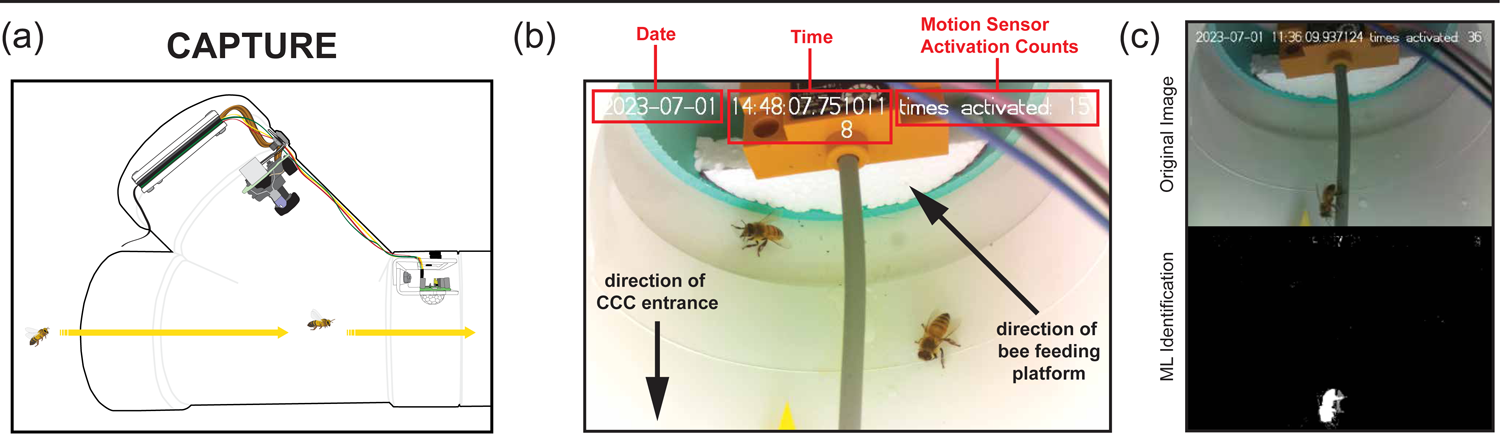
Image and video capture methods. The collection capabilities demonstrated through image and video capture and ML counting methods (a) Cross-section of the apparatus highlighting the image-capture component, demonstrating the configuration responsible for data acquisition.(b) A screenshot of an image captured by the apparatus, displaying metadata at the top, showcasing the comprehensive information gathered during the image-capturing process. (c) A comparison between a RAW image and the same image processed by a ML model designed to identify and annotate bees within the system, illustrating the efficacy of the image analysis component.

### Standalone IR counter performance

Examining the effectiveness of the standalone IR beam counter located at the base of the nectar feeder (**Fig. 3A**) required testing of previously established honeybee preferences for sugary solutions. In this instance, our study compared the counts of bee visits to an apparatus containing water alone to one containing a sugar solution (i.e., artificial nectar). These experiments were carried out over four days, with one experiment being done in the morning (m) before feeder positions were switched in the afternoon (a). In a bulk analysis of four trials with two replicates each (morning and afternoon), bees demonstrated a significant preference for the sugar solution over water alone when comparing the total visits using a two-tailed paired t-test (**Fig. 3B**; p=0.0026). Breaking the total visits down into individual trials also showed a significant difference in bee visitation rates for each day (**Fig. 3C**, trial 1 p=0.0196; all other trials p=<0.0001). The results were consistent with previously established preferences that bees have for sugars(Scheiner et al., 2004; Wykes, 1952), indicating that the apparatus effectively captures bee visits regardless of the position of the food source. The total counts and ratios of artificial nectar-to-water over successive days illustrated the learning behavior of bees. That is, as the bees learned the location of the sugar solution, the signal amplified through the hive, causing dramatic increases in visitation rates over time to the artificial nectar over water alone (**Fig. 3C**).

**Figure 3.**
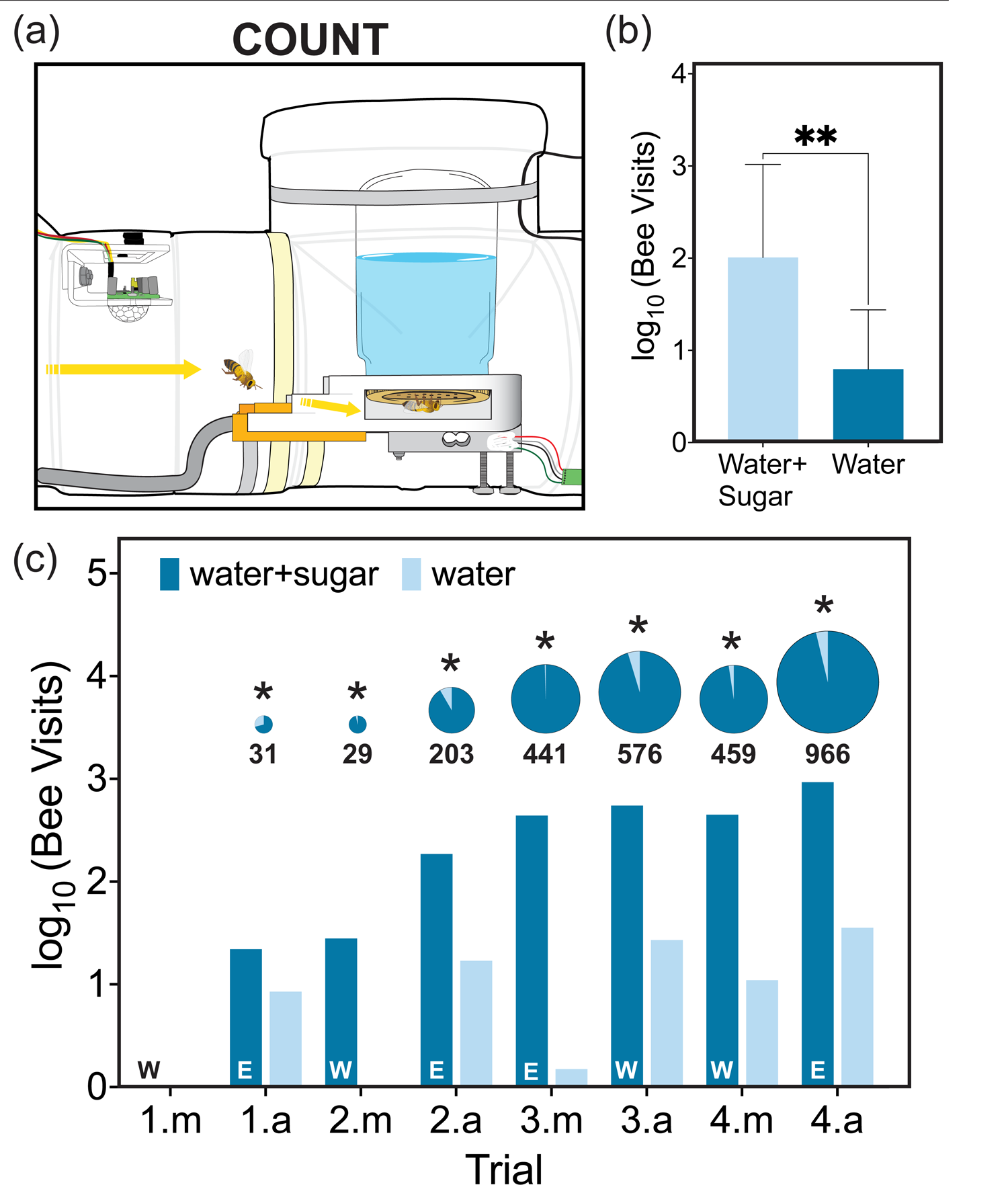
Automated bee counting capability. (a) Cross-section of the apparatus, highlighting the automated standalone IR beam counter. (b) Comparative analysis of total bee visits over a four-day period to apparatuses containing water or artificial nectar (20% sugar solution). (c)Trial-specific results, displaying log-transformed counts of bee visits during each trial, further segmented into morning (trial #.m) and afternoon (trial #.a) sessions. The capacity of the apparatus to monitor changing bee preferences is highlighted as the solutions were switched between morning and afternoon (i.e., the east unit moving to the west mounting position) represented by the E (east) or W (west). Above each trial is a bold number representing the total number of visits between the two apparatuses. At the same time, the pie chart below illustrates the proportion of visits to each apparatus during each trial, with the diameter of the circle representing the total number of visits. All trials with visitations showed a significant difference (*) between the visitation rates of the sugar as compared to the water (trial 1.a p=0.0196; all other trials p=<0.0001; via a Chi-squared test).

An examination of visitation differences between water alone and artificial nectar sheds light on the potential to use the CCC to detect variation in bee preferences over short temporal and spatial spans. Specifically, the apparatus demonstrated the capability of bees to differentiate between reward types even when the entrance to the feeders were located only 50 cm away from one another and when their positions were switched in the middle of the day (also see additional evidence on consumption rates below). This capability may be useful for understanding bee preferences and the factors influencing foraging behavior. The versatility in accommodating various food types also enhances the potential utility of the CCC apparatus.

### Consumption rate by weight

The IR beam counter appeared to quantify bee visitation to the feeders successfully, but it did not measure consumption. To validate the IR beam counter numbers, consumption rates were simultaneously measured by incorporating data from digital weight scales mounted to the bottom of the feeding platforms (**Fig. 4A**). The resulting quantitative data directly assessed the amount of solution consumed in 30-minute intervals (**Fig. 4B**). Bees consumed ca. 20 to 30 g of artificial nectar in two separate trials, whereas an immeasurable amount was consumed from the feeder containing just water (**Fig. 4B**, trial 3 p=0.0011; trial 4 p=0.0033). These results are entirely consistent with the visitation numbers obtained from the standalone IR counters (**Fig. 3C** and presented again as insets in **Fig. 4B**). That is, bees both visited and consumed more solution from the feeders containing artificial nectar versus ones containing water alone, which also validated the data from both the IR and weight sensors. One limitation of this approach is that the counts from the IR beam counter (not run by the Raspberry Pi) were only manually recorded at the end of each 3-hour period over the course of each trial. To overcome this limitation, one could either manually record the counts at specified times throughout the course of a given experiment or incorporate a different IR beam system that records data over time (ideally run by the Raspberry Pi).

**Figure 4.**
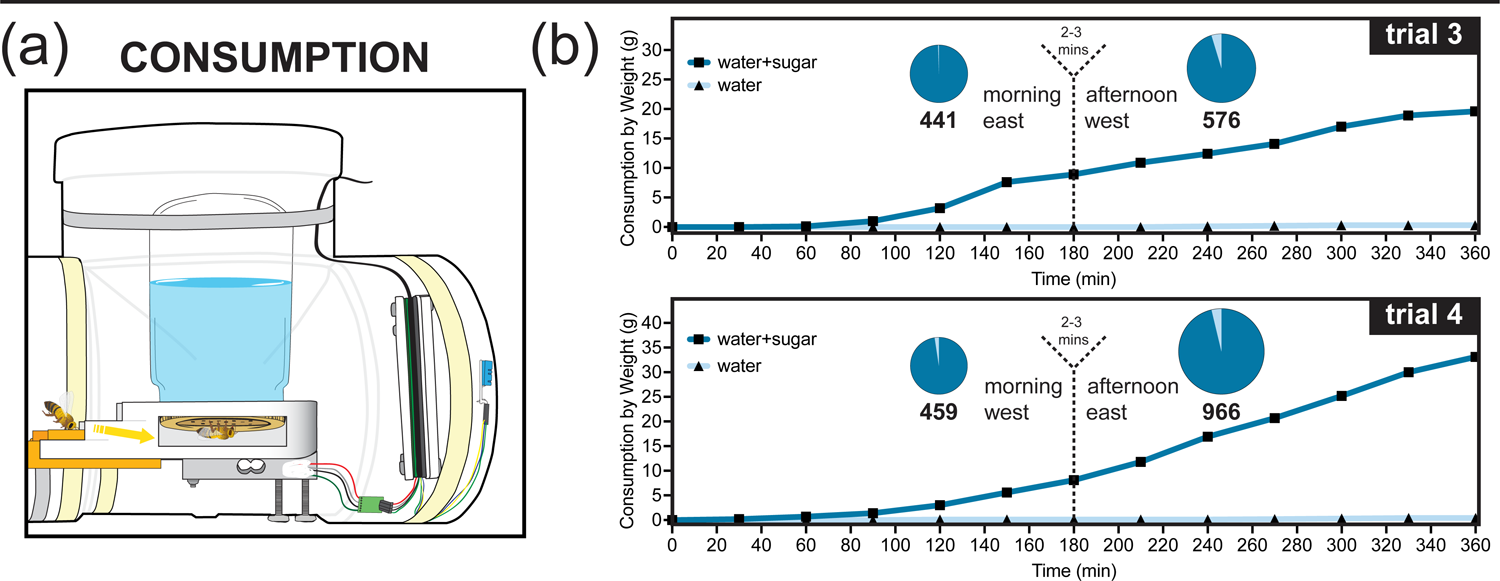
Measure of Consumption. (a) A cross-section of the apparatus, revealing its internal components designed to simultaneously collect consumption rates and environmental data. (b) A comparative analysis demonstrating the consumption rates of artificial nectar (water+sugar) solution versus water alone. The dotted line represents the time at which the position of the traps was switched, with the associated orientation of the artificial nectar noted. The time it took to switch feeder orientation averaged 2-3 minutes, which is noted above the dotted line. A t-test showed a significant difference on both days between the sugar solution and water (trial 3 p=0.0011 and trial 4 p=0.0033).

Interestingly, there was no apparent lag in consumption of the artificial nectar when its position was switched in the middle of a trial (**Fig. 4B**). This result can possibly be attributed to the fact that data from the scales was only recorded every 30 minutes, so any lag in visitation was not recorded before the bees learned the new position of the artificial nectar. As mentioned above, this limitation can be overcome by coding the Raspberry Pi to take more frequent measurements from the scale. At a minimum, it appears that bees are able to rapidly adjust their behavior in response to the location of a reward. It will be interesting to use the CCC apparatus further to tease apart aspects of bee learning and behavior.

### Correlations between environmental conditions and bee activity

Environmental conditions significantly affect bee behavior(Kaehler et al., 2021; Russell & McFrederick, 2021b; Sandoval-Molina et al., 2020b). Temperature and humidity measurements were collected using a dedicated environmental sensor connected to a Raspberry Pi unit **(Table S2** and **D.7 in Fig. S4)**. Measurements were systematically collected every 30 minutes throughout the trial periods. Both temperature and humidity are known to influence bee foraging behavior and may be important factors to consider when investigating bee preferences and foraging activity(Kaehler et al., 2021; Russell & McFrederick, 2021b; Sandoval-Molina et al., 2020b). An analysis of the consumption as a function of temperature indicates a positive correlation between temperature and total nectar consumed (p=<0.001). Exploring the influence of these environmental factors can further contribute to a more comprehensive understanding of bee behavior in certain conditions. Understanding these correlations has implications for modeling and managing bee activities in response to changing environmental conditions, with potential application in agriculture and conservation efforts.

### Limitations of the CCC apparatus

While the CCC apparatus appears useful for assessing bee visitation, behavior, and reward consumption, several important limitations need to be addressed. For instance, we are currently unable to collect individual-level data, but the potential to do so exists with the refinement of ML approaches. Similarly, this system, as used, is unlikely to be able to discern innate (gustatory) from learned (post-ingestive) responses to rewards. The experimental setup also has potential issues with pseudoreplication since a single set of feeders in a single location was used. In an ideal situation, multiple feeders over multiple sites and hives would be used. Of course, the CCC apparatus as designed does not reflect a natural system and currently ignores the broader context provided by flowers, such as colors, patterns, shapes, and scents; however, these aspects could readily be added given the design modularity (**Fig. S2**). Additional technical improvements include refining the sensitivity of the PIR sensor to allow for better direct counting, automating the count and identification of pollinators using AI, placing the camera in closer proximity to the entrance of the feeding platform, decreasing the amount of time between weight measurements, placing a camera or PIR inside of the feeding chamber, and expanding the range of the parameters measured for a more comprehensive analysis.

### Implications and applications

The CCC Apparatus has practical applications in hive monitoring, bee behavioral research, and ecological and agricultural studies. The results from this study and the development of the CCC Apparatus contribute to the fields of bee research by providing an inexpensive, efficient, reliable, and modular apparatus for accurately and automatically counting bees and assessing their preferences. The results from our pilot study demonstrate the efficacy of the system, which provides researchers and beekeepers with an automated tool for monitoring populations and investigating preferences. The insights gained in monitoring bee foraging behaviors can enhance our understanding of bee behavior in natural and agricultural settings, but the system could easily be applied to greenhouse or laboratory settings.

Outside of bee-specific research, this modular platform opens up possibilities for adaptation in studying the foraging behavior of other insects, small birds, mammals, or even reptiles. By modifying the feeding platform, sensors, and system openings, the apparatus can provide valuable insights into the foraging preferences and ecological interactions of various species. The adaptability and modularity of the design and integrated components represent an avenue for exploring even remote locations, providing opportunities to study rare and endangered species remotely and cost-effectively.

## ACKNOWLEDGEMENTS

This work was supported by a grant from the US NSF to ECS-R, ADH, and CJC. We thank the University of Minnesota Bee Lab and Drs. Marla Spivak and Katie Lee for helping to develop and test the apparatus while simultaneously allowing us to use their honeybee populations for the purpose of this study. We also thank Dr. Neil E. Olszewski and Calvin Peters from the University of Minnesota for providing materials and additional research support, including the Raspberry Pi and camera systems, as well as consulting on pollinator behavior and A.I. development.

## AUTHOR CONTRIBUTIONS

E.M., J.N., E.S.R., A.H., and CC conceived the experimental approach. E.M. and CC conducted the experiments. E.M. performed the literature review. J.N. and E.M. wrote the code for the computational components of the apparatus. E.M. collected and analyzed the data. E.M. wrote a majority of the manuscript with contributions from all co-authors.

## SUPPORTING INFORMATION

**Fig. S1** – Design and assembly of the CCC Apparatus

**Fig. S2** – Possible alternative CCC designs based on modularity

**Fig. S3** – Experimental setup for testing the CCC Apparatus

**Fig. S4** – Correlation of temperature to nectar consumption

**Table S1** – Overview of methods for monitoring bees

**Table S2** – Parts used in the assembly of the CCC

## REFERENCES

Agatz, A., Miles, M., Roeben, V., Schad, T., Stouwe, F., Zakharova, L., & Preuss, T. G. (2023). Evaluating and Explaining the Variability of Honey Bee Field Studies across Europe Using BEEHAVE. Environmental Toxicology and Chemistry, 42(8), 1839– 1850. 10.1002/etc.5678

Alburaki, M., Madella, S., & Corona, M. (2021). RFID Technology Serving Honey Bee Research: A Comprehensive Description of a 32-Antenna System to Study Honey Bee and Queen Behavior. Applied System Innovation, 4(4), 88. 10.3390/asi4040088

Alqarni, A. S., Ali, H., Iqbal, J., & Raweh, H. S. A. (2023). Proboscis Extension Response of Three Apis mellifera Subspecies toward Water and Sugars in Subtropical Ecosystem. Stresses, 3(1), 182–197. 10.3390/stresses3010014

Anderson, C. (2023). Pollen Feeder for Bees-Make Your Own. Carolina Honeybees. https://carolinahoneybees.com/make-a-pollen-feeder/

Atakan, E., & Pehlivan, S. (2015). Attractiveness of various colored sticky traps to some pollinating insects in apple. TURKISH JOURNAL OF ZOOLOGY, 39, 474–481. 10.3906/zoo-1403-62

Avisar, D., Azulay, S., Bombonato, L., Carvalho, D., Dallapicolla, H., de Souza, C., dos Santos, A., Dias, T., Galan, M. P., Galvao, M., Gonsalves, J. M., Gonzales, E., Graça, R., Livne, S., Mafia, R., Manoeli, A., May, M., Menezes, T. R. D., Pinheiro, A. C., … Silva, W. (2023). Safety Assessment of the CP4 EPSPS and NPTII Proteins in Eucalyptus. GM Crops & Food, 14(1), 1–14. 10.1080/21645698.2023.2222436

Balfour, N. J., & Ratnieks, F. L. W. (2023). Why Petals? Naïve, but Not Experienced Bees, Preferentially Visit Flowers with Larger Visual Signals. Insects, 14(2), 130. 10.3390/insects14020130

Campbell, A. J., Gomes, R. L. C., da Silva, K. C., & Contrera, F. A. L. (2019). Temporal variation in homing ability of the neotropical stingless bee Scaptotrigona aff. postica (Hymenoptera: Apidae: Meliponini). Apidologie, 50(5), 720–732. 10.1007/s13592-019-00682-z

Corbet, S. A., Fussell, M., Ake, R., Fraser, A., Gunson, C., Savage, A., & Smith, K. (1993). Temperature and the pollinating activity of social bees. Ecological Entomology, 18(1), 17–30. 10.1111/j.1365-2311.1993.tb01075.x

Dainese, M., Martin, E. A., Aizen, M. A., Albrecht, M., Bartomeus, I., Bommarco, R., Carvalheiro, L. G., Chaplin-Kramer, R., Gagic, V., Garibaldi, L. A., Ghazoul, J., Grab, H., Jonsson, M., Karp, D. S., Kennedy, C. M., Kleijn, D., Kremen, C., Landis, D. A., Letourneau, D. K., … Steffan-Dewenter, I. (2019). A global synthesis reveals biodiversity-mediated benefits for crop production. Science Advances, 5(10), eaax0121. 10.1126/sciadv.aax0121

Descamps, C., Quinet, M., Baijot, A., & Jacquemart, A. (2018). Temperature and water stress affect plant–pollinator interactions in Borago officinalis (Boraginaceae). Ecology and Evolution, 8(6), 3443–3456. 10.1002/ece3.3914

Fernandes, N., Luz, L., Filho, E. A., de Aragão, F. A., Zocolo, G., & Freitas, B. (2023). Differences in the Chemical Composition of Melon (Cucumis melo L.) Nectar Explain Flower Gender Preference by Its Pollinator, Apis mellifera. Journal of the Brazilian Chemical Society. 10.21577/0103-5053.20230010

Garibaldi, L. A., Steffan-Dewenter, I., Winfree, R., Aizen, M. A., Bommarco, R., Cunningham, S. A., Kremen, C., Carvalheiro, L. G., Harder, L. D., Afik, O., Bartomeus, I., Benjamin, F., Boreux, V., Cariveau, D., Chacoff, N. P., Dudenhöffer, J. H., Freitas, B. M., Ghazoul, J., Greenleaf, S., … Klein, A. M. (2013). Wild Pollinators Enhance Fruit Set of Crops Regardless of Honey Bee Abundance. Science, 339(6127), 1608–1611. 10.1126/science.1230200

Gernat, T., Jagla, T., Jones, B. M., Middendorf, M., & Robinson, G. E. (2023). Automated monitoring of honey bees with barcodes and artificial intelligence reveals two distinct social networks from a single affiliative behavior. Scientific Reports, 13(1), 1541. 10.1038/s41598-022-26825-4

Harmon-Threatt, A. N., & Anderson, N. L. (2023). Bee movement between natural fragments is rare despite differences in species, patch, and matrix variables. Landscape Ecology, 38(10), 2519–2531. 10.1007/s10980-023-01719-6

Kaehler, T. G., Halinski, R., Contrera, F. A. L., Silveira, A., & Blochtein, B. (2021). Flight distance and foraging of Tetragonisca fiebrigi (Apidae: Meliponini) in response to different concentrations of sugar in food resources and abiotic factors. Journal of Apicultural Research, 1–13. 10.1080/00218839.2021.2005872

Layek, U., Das, N., Kundu, A., & Karmakar, P. (2022). Methods Employed in the Determining Nectar and Pollen Sources for Bees: A Review of the Global Scenario. Annals of the Entomological Society of America, 115(6), 417–426. 10.1093/aesa/saac013

Leach, M., Dibble, A. C., Stack, L. B., Perkins, L. B., & Drummond, F. A. (2023). The Effect of Plant Nutrition on Bee Flower Visitation. Journal of the Kansas Entomological Society, 94(4), 277–300. 10.2317/0022-8567-94.4.277

Lehrer, M., Horridge, G. A., Zhang, S. W., & Gadagkar, R. (1995). Shape vision in bees: innate preference for flower-like patterns. Philosophical Transactions of the Royal Society of London. Series B: Biological Sciences, 347(1320), 123–137. 10.1098/rstb.1995.0017

Lemanski, N. J., Williams, N. M., & Winfree, R. (2022). Greater bee diversity is needed to maintain crop pollination over time. Nature Ecology & Evolution, 6(10), 1516–1523. 10.1038/s41559-022-01847-3

Liao, C., Xu, Y., Sun, Y., Lehnert, M. S., Xiang, W., Wu, J., & Wu, Z. (2020). Feeding behavior of honey bees on dry sugar. Journal of Insect Physiology, 124, 104059. 10.1016/j.jinsphys.2020.104059

Liira, J., & Jürjendal, I. (2023). Are bees attracted by flower richness? Implications for ecosystem service-based policy. Ecological Indicators, 154, 110927. 10.1016/j.ecolind.2023.110927

Lowenstein, D. M., Matteson, K. C., & Minor, E. S. (2015). Diversity of wild bees supports pollination services in an urbanized landscape. Oecologia, 179(3), 811–821. 10.1007/s00442-015-3389-0

Lüthi, M. N., Berardi, A. E., Mandel, T., Freitas, L. B., & Kuhlemeier, C. (2022). Single gene mutation in a plant MYB transcription factor causes a major shift in pollinator preference. Current Biology, 32(24), 5295–5308.e5. 10.1016/j.cub.2022.11.006

Ma, W., Long, D., Wang, Y., Li, X., Huang, J., Shen, J., Su, W., Jiang, Y., & Li, J. (2021). Electrophysiological and behavioral responses of Asian and European honeybees to pear flower volatiles. Journal of Asia-Pacific Entomology, 24(1), 221–228. 10.1016/j.aspen.2020.12.011

Mayack, C., Cook, S. E., Niño, B. D., Rivera, L., Niño, E. L., & Seshadri, A. (2023). Poor Air Quality Is Linked to Stress in Honeybees and Can Be Compounded by the Presence of Disease. Insects, 14(8), 689. 10.3390/insects14080689

McCravy, K. W., & Ruholl, J. D. (2017). Bee (Hymenoptera: Apoidea) Diversity and Sampling Methodology in a Midwestern USA Deciduous Forest. Insects, 8(3), 81. 10.3390/insects8030081

Murphree, S. M. (2022). Effects of imidacloprid and octopamine on the honey bee (Apis mellifera) trophallaxis social network.

Mustard, J. A., Oquita, R., Garza, P., & Stoker, A. (2019). Honey Bees (Apis mellifera) Show a Preference for the Consumption of Ethanol. Alcoholism: Clinical and Experimental Research, 43(1), 26–35. 10.1111/acer.13908

Muth, F., Cooper, T. R., Bonilla, R. F., & Leonard, A. S. (2018). A novel protocol for studying bee cognition in the wild. Methods in Ecology and Evolution, 9(1), 78–87. 10.1111/2041-210x.12852

Nicolson, S. W. (2022). Sweet solutions: nectar chemistry and quality. Philosophical Transactions of the Royal Society B, 377(1853), 20210163. 10.1098/rstb.2021.0163

Nouvian, M., & Galizia, C. G. (2019). Aversive Training of Honey Bees in an Automated Y-Maze. Frontiers in Physiology, 10, 678. 10.3389/fphys.2019.00678

Odemer, R. (2022). Approaches, challenges and recent advances in automated bee counting devices: A review. Annals of Applied Biology, 180(1), 73–89. 10.1111/aab.12727

Papa, G., Maier, R., Durazzo, A., Lucarini, M., Karabagias, I. K., Plutino, M., Bianchetto, E., Aromolo, R., Pignatti, G., Ambrogio, A., Pellecchia, M., & Negri, I. (2022). The Honey Bee Apis mellifera: An Insect at the Interface between Human and Ecosystem Health. Biology, 11(2), 233. 10.3390/biology11020233

Pešović, U., Ranđić, S., & Stamenkovic, Z. (2017). *Design and Implementation of Hardware Platform for Monitoring Honeybee Activity*.

Prasad, A. V., & Hodge, S. (2013). Factors influencing the foraging activity of the allodapine bee Braunsapis puangensis on creeping daisy (Sphagneticola trilobata) in Fiji. Journal of Hymenoptera Research, 35(35), 56–69. 10.3897/jhr.35.6006

Prendergast, K. S., Menz, M. H. M., Dixon, K. W., & Bateman, P. W. (2020). The relative performance of sampling methods for native bees: an empirical test and review of the literature. Ecosphere, 11(5). 10.1002/ecs2.3076

Richman, S. K., Muth, F., & Leonard, A. S. (2021). Measuring foraging preferences in bumble bees: a comparison of popular laboratory methods and a test for sucrose preferences following neonicotinoid exposure. Oecologia, 196(4), 963–976. 10.1007/s00442-021-04979-8

Russell, K. A., & McFrederick, Q. S. (2021a). Elevated Temperature May Affect Nectar Microbes, Nectar Sugars, and Bumble Bee Foraging Preference. Microbial Ecology, 1–10. 10.1007/s00248-021-01881-x

Russell, K. A., & McFrederick, Q. S. (2021b). Elevated Temperature May Affect Nectar Microbes, Nectar Sugars, and Bumble Bee Foraging Preference. Microbial Ecology, 1–10. 10.1007/s00248-021-01881-x

Sandoval-Molina, M. A., Flórez-Gómez, N. A., Pérez-Botello, A. M., Hinojosa-Díaz, I. A., Reyes-Tovar, J. M., & Ayala, R. (2020a). Effects of floral display and abiotic environment on the foraging activity of bees on Kallstroemia pubescens (Zygophyllaceae). Ethology Ecology & Evolution, 32(6), 551–571. 10.1080/03949370.2020.1755371

Scheiner, R., Page, R. E., & Erber, J. (2004). Sucrose responsiveness and behavioral plasticity in honey bees (Apis mellifera). Apidologie, 35(2), 133–142. 10.1051/apido:2004001

Sharif, M. Z., Di, N., & Liu, F. (2022). Monitoring honeybees (Apis spp.) (Hymenoptera: Apidae) in climate-smart agriculture: A review. Applied Entomology and Zoology, 57(4), 289–303. 10.1007/s13355-021-00765-3

Simpson, D. T., Weinman, L. R., Genung, M. A., Roswell, M., MacLeod, M., & Winfree, R. (2022). Many bee species, including rare species, are important for function of entire plant–pollinator networks. Proceedings of the Royal Society B, 289(1972), 20212689. 10.1098/rspb.2021.2689

Skvarla, M. J., Larson, J. L., Fisher, J. R., & Dowling, A. P. G. (2020). A Review of Terrestrial and Canopy Malaise Traps. Annals of the Entomological Society of America, 114(1), 27–47. 10.1093/aesa/saaa044

Su, W., Ma, W., Zhang, Q., Hu, X., Ding, G., Jiang, Y., & Huang, J. (2022). Honey Bee Foraging Decisions Influenced by Pear Volatiles. Agriculture, 12(8), 1074. 10.3390/agriculture12081074

Tong, Z.-Y., Wu, L.-Y., Feng, H.-H., Zhang, M., Armbruster, W. S., Renner, S. S., & Huang, S.-Q. (2023). New calculations imply that 90% of flowering plant species are animal-pollinated. National Science Review. 10.1093/nsr/nwad219

Vaudo, A. D., Biddinger, D. J., Sickel, W., Keller, A., & López-Uribe, M. M. (2020). Introduced bees (Osmia cornifrons) collect pollen from both coevolved and novel host-plant species within their family-level phylogenetic preferences. Royal Society Open Science, 7(7), 200225. 10.1098/rsos.200225

Visscher, P., & Seeley, T. (2023). *Bee-Lining as a Research Technique in Ecological Studies of Honey Bees*.

Wagner, D. L. (2019). Insect Declines in the Anthropocene. Annual Review of Entomology, 65(1), 1–24. 10.1146/annurev-ento-011019-025151

Williams, S. M., Aldabashi, N., Cross, P., & Palego, C. (2023). Challenges in Developing a Real-Time Bee-Counting Radar. Sensors, 23(11), 5250. 10.3390/s23115250

Wykes, G. R. (1952). The Preferences of Honeybees for Solutions of Various Sugars Which Occur in Nectar. Journal of Experimental Biology, 29(4), 511–519. 10.1242/jeb.29.4.511

