## Supplemental Tables and Figures for "Methods for assessing bee foraging preferences: a short review and a new automated apparatus"

Supplemental Table 1. Methods for Sampling and Testing Preferences of Apis spp. Overview of the methodologies employed in sampling bee populations and assessing their preferences, offering insights into their distinct applications, data collecting procedures, costs, and other pertinent aspects. These methods are then compared to the capabilities of the CCC apparatus, highlighting the unique advantages it offers for bee research and monitoring.

| Method | Source | System Overview | Data Collection | Suitable Setting | Cost | Labor | Stress Inducing | Visitation Rate | Visit Duration | Data Storage | Consumption Rate | Environmental Conditions | Species Identification | Foraging Behavior | Nectar Collection | Pollen Collection | Image/Video Collection | Colony Level Data | All Weather |
|---|---|---|---|---|---|---|---|---|---|---|---|---|---|---|---|---|---|---|---|
| Visual Observation | Leach et al. 2023 | researchers observe and record insect visitations to specific plants or areas over a period of time, and associated behaviors | direct | field | low | high | - | ● | ○ | | ○ | ● | ○ | ● | | | ○ | ● | ○ |
| Sweep Netting | MG et al. 2023 | nets are used to collect insects within in environment or plant canopy | direct | field | low | high | +/- | | | | | | ○ | | | ● | | | |
| Malaise Traps | Skvarla et al. 2021 | passive traps that use a barrier of fabric or netting to direct flying insects into a collecting container attached to the trap | direct | field | low | high | +/- | | | | | | ○ | | | | | | |
| Sticky Traps | Atakan et al. 2015 | sheets or cards coated with a sticky substance that insects adhere to when they come into contact with it | direct | field | low | high | + | | | | | | ○ | | | | | | |
| Windowpane Traps | Visscher and Seeley 1989 | specialized traps that intercept flying insects in flight | direct | field | low | high | +/- | | | | | | ○ | | | | | | |
| Pan traps/bowl traps/bee bowls/ Moericke traps | McCravy and Ruholl 2017 | shallow containers filled with a liquid that attracts insects and fall into the trap | direct | field | low | high | +/- | | | | | | ○ | | | | | | |
| Feeding Platforms with Cameras | Murphree 2022 | platforms are equipped with cameras that capture video footage of the feeding activity | remote | field, free-flight enclosures | high | low | - | ● | ○ | ● | ○ | | ○ | ● | ○ | ● | ○ | | |
| Mobile Gardens | Lowenstein et al. 2015 | researchers create portable gardens with specific plants to attract and observe insects, enabling the study of their visitation patterns | direct | field | low | high | - | ● | ○ | | | | ○ | ● | ○ | | | | |
| Video Recordings and Analysis | Gernat 2023 | cameras and software (AI) are utilized to analyze the footage and accurately count the number of bee or insect visits | remote | field, free-flight enclosures, laboratory | high | low | - | ● | ○ | ● | | | ○ | ● | | | ○ | | |
| Beating Sheets | Avisar et al. 2023 | sheet spread beneath a plant or shrub,which is then gently beaten or shaken, causing the insects to fall onto the sheet | direct | field | low | high | +/- | | | | | | ○ | | | | | | |
| Acoustic Monitoring | Sharif et al. 2023 | sensitive equipment, such as accelerometers and microphones, to capture the sounds or vibrations produced by insects for communication or other activities | remote | field | high | high | - | | | ● | | | | | | | | ● | |
| Doplar Radar | Williams et al. 2023 | utilized to count insects by taking advantage of their ability to reflect radar signals | remote | field | high | low | - | | | ● | | ● | | ● | | | | ● | ○ |
| Mark-Recapture Study | Harmon-Threatt and Anderson 2023 | bees are marked with paint and released into the wild, recapturing and identiying tagged bees informs floral preferences | direct | field | low/high | high | +/- | ● | ○ | | | | ○ | | | | | ● | |
| Pollen Analysis | Layek et al. 2022 | preferences can be determined by pollen loads on their bodies and in their corbiculae, detect pollen can be traced back to flowers visited | direct | field | high | high | +/- | ● | | | | | | | | ● | | | ○ |
| Choice Experiment | Mustard et al. 2018 | bees presented with different choices ranging from volatiles to colors and are allowed to choose, can be used in field and laboratory settings | direct / remote | field, free-flight enclosures, laboratory | low / high | high | +/- | ● | ○ | | ○ | | | ● | ○ | ● | | | ○ |
| Electrophysiological Recordings | Ma et al. 2021 | researchers measure the neural responses of bees when exposed to different sensory cues, such as flower scents or colors | direct | laboratory | high | high | + | | | | ○ | | | | | | | | ○ |
| Y-Maze or T-Maze Test | Nouvian and Galizia 2019 | laboratory setups involve offering bees a choice between two or more options to determine their preferences for certain odors, colors, or other stimuli | direct | laboratory | low/high | high | - | ● | ○ | | ○ | | | | ○ | | | | ○ |
| Chemical Analysis | Fernandes et al. 2023 | analyze the chemical composition of flower nectar and pollen to identify compounds that might attract bees | direct | laboratory | high | high | - | | | | | | | | | | | | ○ |
| Radio Frequency Identification (RFID) Tracking | Alburaki et al. 2021 | RFID technology to track the movements of individual bees within a controlled environment | remote | field, free-flight enclosure | high | high | +/- | ● | ○ | | | | | ● | ○ | | | ● | |
| Genetic and Molecular Techniques | Lüthi et al. 2022 | researchers study th genes and molecular pathways involved in preference, such as genes associated with color perception | direct | field, laboratory | high | high | +/- | | | | | | ○ | | | | | ● | |
| Phylogenetic Analysis | Vaudo et al. 2020 | comparative study across bee species to determine patterns and preferences among different lineages | computational | laboratory | low/high | high | +/- | | | | | | ○ | | | | | | ○ |
| Mathematical Modeling | Liao et al. 2020 | models used to simulate and predict bee foraging behavior based on various factors | computational | laboratory | low/high | high | - | | | ● | | | | | | | | ● | ○ |
| Proboscis Extension Reflex (PER) Assay | Alqarni et al. 2023 | laboratory test in which bees extend their proboscis in reaction to odor or stimulii, indicating preference | direct | laboratory | high | high | +/- | | | | ○ | | | | ○ | | | | ○ |
| Free Moving Proboscis Extension Response (FMPER) Assays | Muth et al. 2018 | adaptation of PER assay to measure the color preferences, learning performance and memory of wild bees | direct | field, free-flight enclosure | low/high | high | - | ● | ○ | | ○ | | | ● | ○ | | | | |
| Restricted Volume Preference Assay (RVPA) | Richman et al. 2020 | small enclosures provide free-movent within the space to choose from small volumes of liquid or gas | remote | laboratory | low/high | high | +/- | ● | ○ | | ○ | | | | ○ | ● | | | |
| Infrared (IR) Sensor | Pešović et al. 2017 | sensors can detect the heat signature of the bees, monitoring motion and counts as bees pass through IR beam | remote | field, free-flight enclosure, laboratory | low/high | low | - | ● | ○ | ● | | | | | | | | ● | |
| Capture, Count, and Consumption (CCC) Apparatus | | sensors can detect and count bee visitations while cameras capture their movement and behavior, additional sensors monitor mass (consumption) and environmental factors | remote | field, free-flight enclosure | low/high | low | - | ● | ○ | ● | ○ | ● | ○ | ● | ○ | ● | ○ | ● | ○ |

Method | Source | System Overview | Data Collection | Suitable Setting | Cost | Labor | Stress Inducing | Visitation Rate | Visit Duration | Data Storage | Consumption Rate | Environmental Conditions | Species Identification | Foraging Behavior | Nectar Collection | Pollen Collection | Image/Video Collection | Colony Level Data | All Weather

Supplemental Table 2. CCC Apparatus Parts List for One Apparatus. The following table presents the components of a singular CCC apparatus. The apparatus consists of several components, each serving a specific purpose.The combination of these components creates an effective tool for studying bee populations and behavior. Each component is accompanied by its respective quantity, function, and placement within the system. In instances where relevant, the manufacturer and part number are also included for reference.

| Part | Manufacturer | Part Number | Quantity | Location in Apparatus | Function |
|---|---|---|---|---|---|
| 4" x 2' Solid PVC Sewer/Drain Pipe (A.2)<br>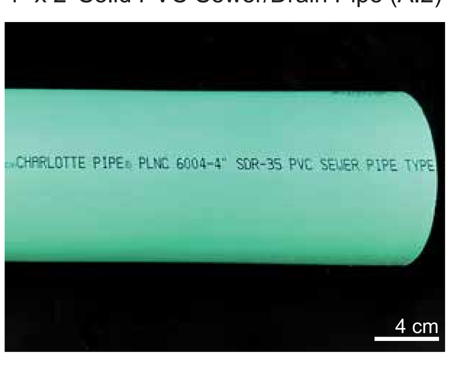 | Charlotte Pipe® | 3034 | **1** | connecting the tee and wye PVC pipes | connecting the tee and wye pipes to one another, while also housing the styrofoam support for the IR beam counter |
| 4" PVC Sewer and Drain Tee (A.3)<br>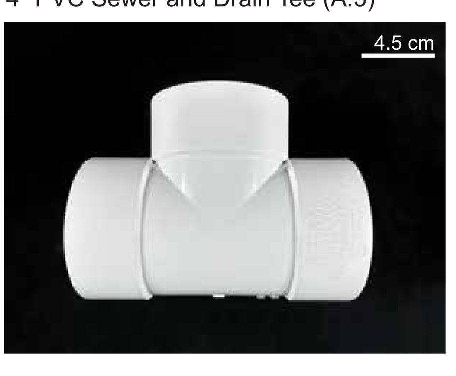 | Tigre® | 36-1084 | **1** | closed end of the system | housing unit at the rear of the apparatus, housing the feeder, weight scale, envirnomental sensor, and Raspberry Pi unit |
| 4” PVC 45 Degree Wye (A.1)<br>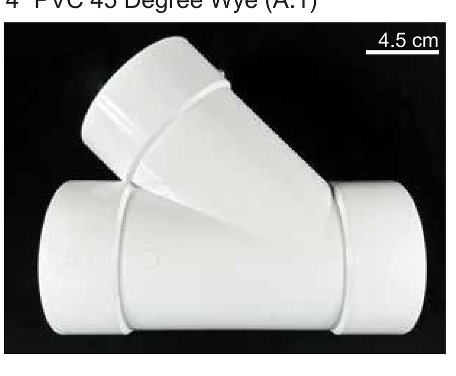 | Tigre® | 36-730 | **1** | open end of the systen | housing unti at the opening of the apparatus, housing the camera, motion detector, and Raspberry Pi |
| 4" PVC Test Cap (A.4)<br>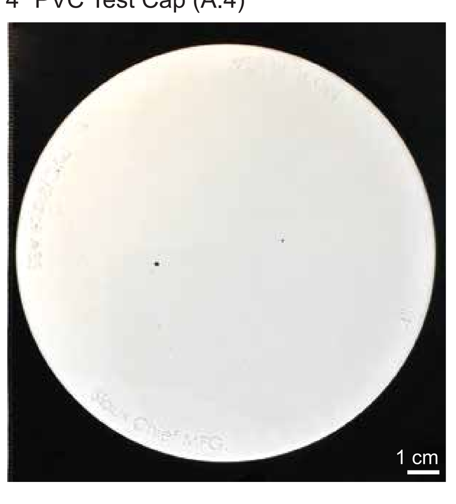 | Sioux Chief® HardHat™ | 880-02PPK | **2** | one placed atop the opening of the wye PVC and one placed atop the opening of the tee pipe | cover the top opening of the wye pipe to shield interior and mount for Rasperry Pi; on the top of the tee pipe to shield interior from biotic and abiotic intrusion |
| 4" PVC Sewer and Drain Grate (A.5)<br>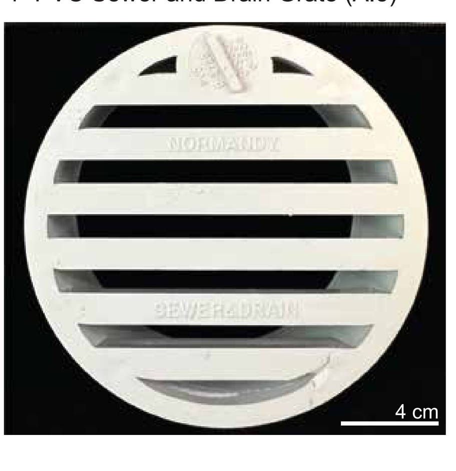 | Tigre® | 36-804 | **1** | inside the tee pipe at the rear of the apparatus | provide airflow through apparatus while preventing insects from entering the rear of the apparatus |
| 1.5” Stainless Steel L-Bracket (B.8)<br>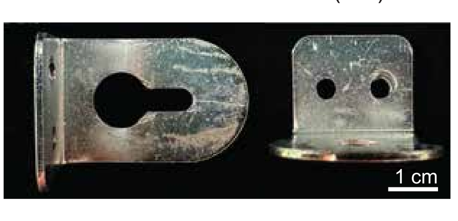 | | | **1** | top portion of the wye pipe, fastened to the ceiling | mounting plate for the Pi camera |
| 10 mm x 3 mm Neodymium Magnets (B.7)<br>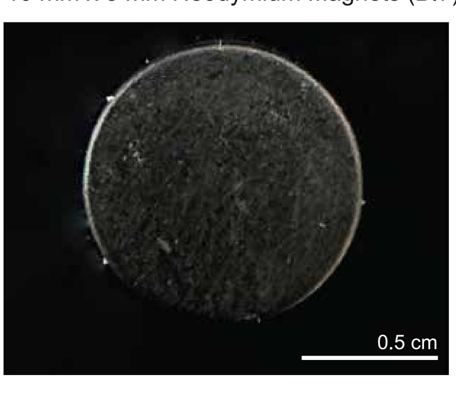 | Nazzo® | | **4** | one set fastened to the ceiling of the horizontal portion of the wye pipe, close to the split, and one fastened to the motion sensor housing | used to mount the motion sensor housing unit to the interior ceiling of the wye pipe |
| 1.9 cm Sticky Back Velcro Tape (B.9)<br>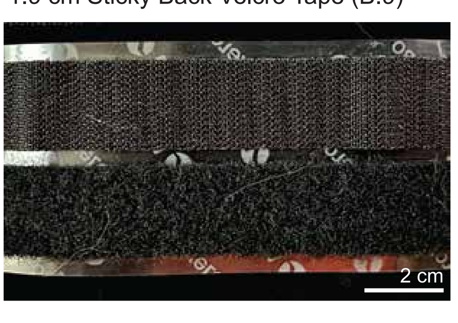 | Velcro® | RN77881 | **~ 40 cm** | backs of Raspberry Pi computers, behind Pi cameras, one side of the L-bracket facing towards the interior, back of the rear styrofoam, interior side of the wye end cap | used to mount the Raspberry Pi computers and Pi camera inside the apparatus |
| IR Pyroelectric Infrared Motion Sensor Detector Module (B.3)<br>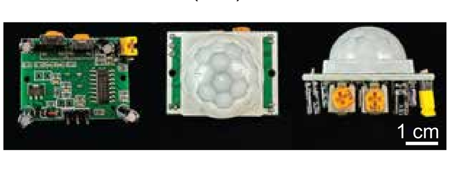 | | HC-SR501 | **1** | inside of a holder, hanging from the ceiling, near the split in the wye pipe | capacity to be used to either count the occurences of motion or as a trigger for the Pi camera |
| Transparent Acrylic Bracket Case Holder for IR Motion Sensor Detector Module (B.4)<br>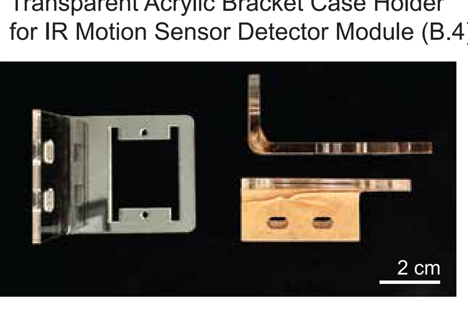 | DORHEA® | | **2** | hanging from the ceiling, near the split in the wye pipe | secure housing for the motion sensor |
| Weight Precision AD Amplifier Breakout Module Pressure Sensor + Load Cell (D.2)<br>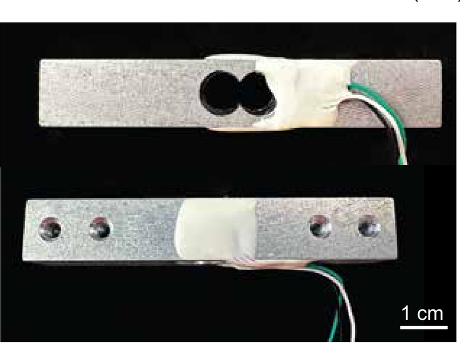 | SazkJere® | HX711 | **1** | inside tee pipe near the middle | measure the mass of the feeder plus the solution, in order to measure consumption |
| DHT11 Temperature Humidity Sensor Module (D.7)<br>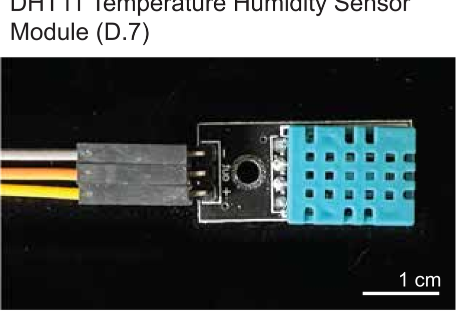 | BOJACK® | DHT11 | **1** | in between the drain grate and foam, with blue block facing outward | measure the temperature and relative humidity of the external or internal environment |
| BERM-6H-30E4-CT Electronic Industrial Sensor Auto Induction IR Beam Count (C.1)<br>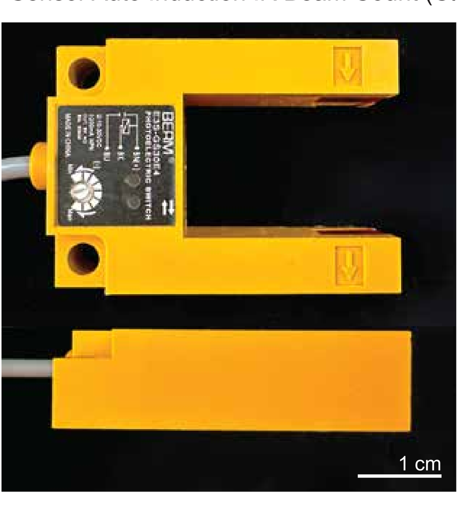 | BERM® | E3S-GS30E4 | **1** | inside of the pipe, right at the opening the beehive feeder | count the passing of bees into and out of the beehive entrance feeding platform |
| Camera Module Auto IR-Cut Switching Day/Night Vision 1080p Camera (B.2)<br>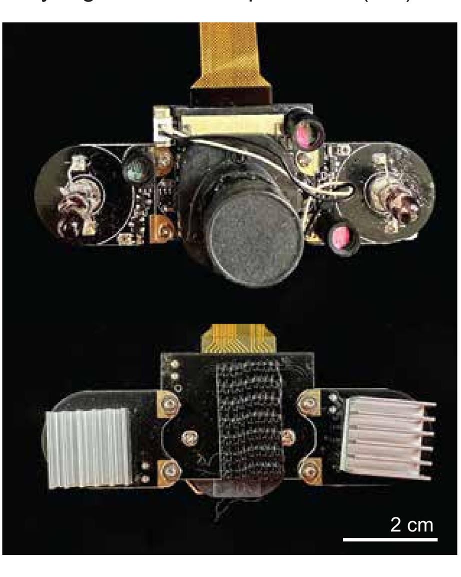 | Longruner | | **1** | upper section of the wye pipe, velcroed to the L-bracket | used to capture images or video inside of the apparatus |
| Raspberry Pi Zero with Transparent Acrylic Protective Case (B.1)<br>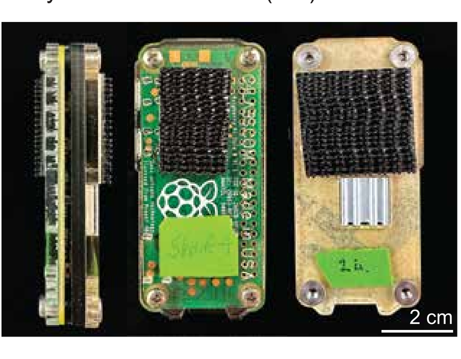 | Raspberry® | | **2** | velcroed to the cap in the top of the wye pipe and velcroed to the foam in the rear of the tee pipe | functions as the primary computing power for the automated counting, image capturing, changes in mass, and recording environmental conditions |
| Plastic Beehive Entrance Feeder (D.1)<br>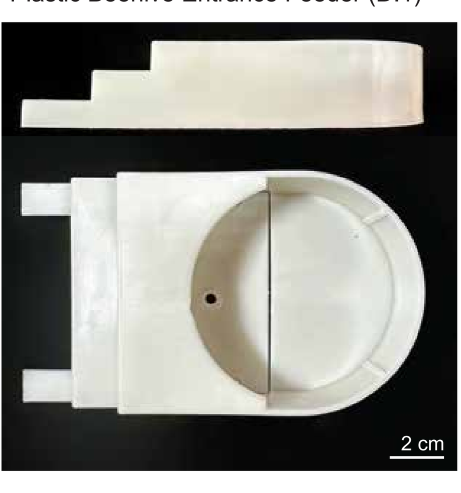 | | | **1** | middle of tee pipe | feeding platform used to lure organisms or to test preferences between food sources |
| Large Plastic Washer (D.3)<br>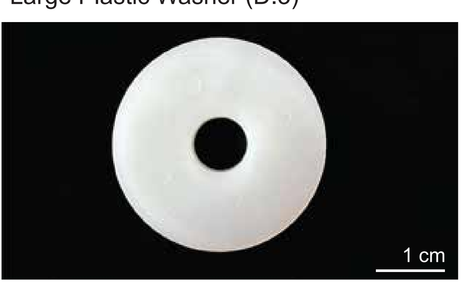 | | | **1** | bottom of feeder, between the feeding platform and weight scale | even weight distribution of the feeding platform upon the weight scale |
| White Plastic Scew Head End Cap (D.4)<br>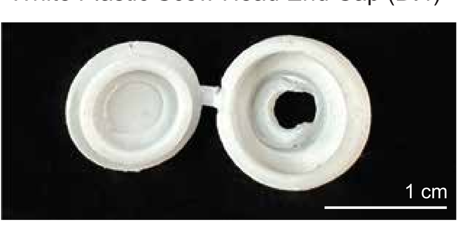 | | | **1** | interior of the feeding platform | protective cap to protect the bees and the connection between the feeding platform and scale |
| 3 cm No. 4 Pan Head Machine Screw and Lock Nut (D.5)<br>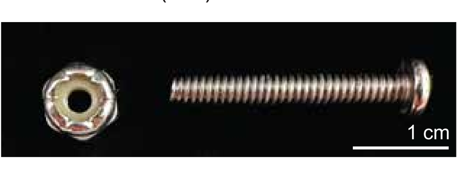 | | | **1 sets** | through the weight scale and feeding platform, inside of the tee pipe | connects the beehive entrance feeder to the weight scale |
| 3 cm No. 5 Oval Head Machine Screw and Lock Nut (D.6)<br>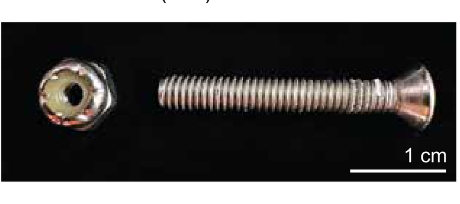 | | | **2 sets** | through the bottom of the tee pipe and into weight scale | suspends the weight scale above the floor of the tee pipe |
| 1.5 cm No. 5 Pan Head Machine Screw, Washer, and Lock Nut (B.5)<br>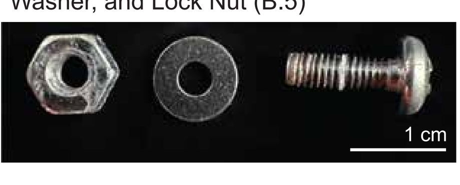 | | | **4 set** | in the housing of the motion sensor, and in the L-bracket | connects the L-bracket to the ceiling of the top portion of the wye pipe and connects the two motion sensor housing units |
| 7 mm M1.2 Oval Head Screw, Washer. and Nut (B.6)<br>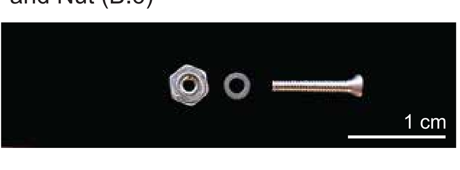 | | | **2 sets** | motion sensor housing | secures the motion sensor inside of the protective housing |
| 1.5 in Open Cell Foam Rubber 10.5 in x 12 in (E.1-E.4)<br>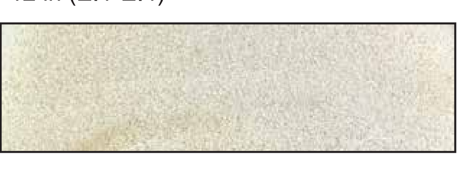 | | | **1** | middle of the tee pipe | provide ventilation through the rear of the apparatus and block organisms from entering the back section of the apparatus, guiding the individuals into the feeding platform |
| 0.5 in Styrofoam 12 in x 6 in (E.5)<br>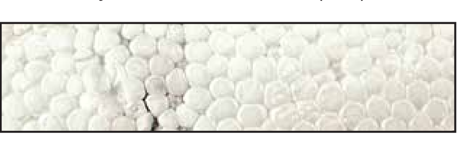 | | | **1** | connector piece between wye and tee pipe | used as support for the back end of the IR beam counter and as a support to keep the feeder tank upright, when sitting on the feeding platform |
| Berm Digital Display Counter (C.2)<br>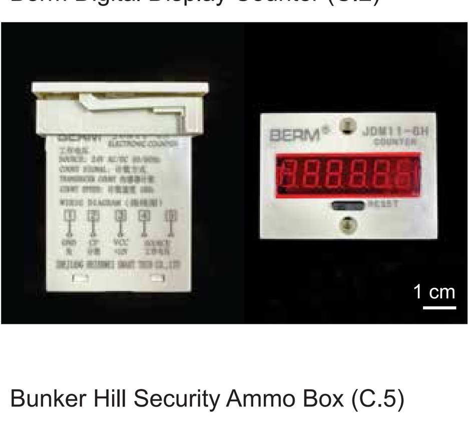 | BERM® | JDM11-6H | **1** | digital read out of IR beam counter | mounted on the side of the ammo box |
| Bunker Hill Security Ammo Box (C.5)<br>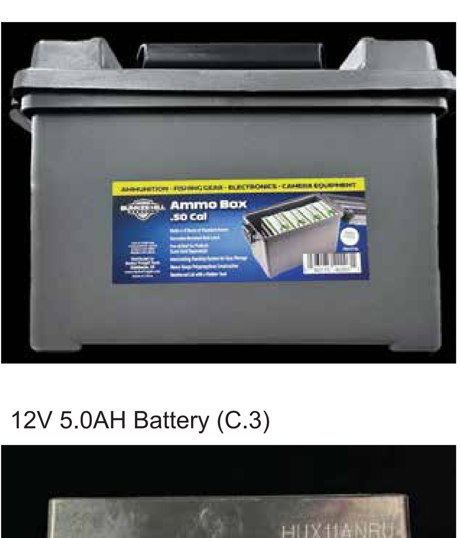 | Bunker Hill Security® | 57766 | **1** | located near the appartus | houses the batteries and houses the counters associated with the IR beam counters |
| 12V 5.0AH Battery (C.3)<br>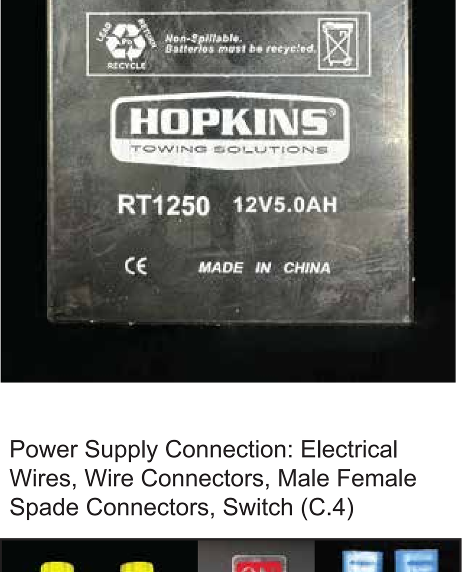 | Hopkins Towing Solutions® | | **1** | inside ammo box | provides power for the IR beam counter and other electronics |
| Power Supply Connection: Electrical Wires, Wire Connectors, Male Female Spade Connectors, Switch (C.4)<br>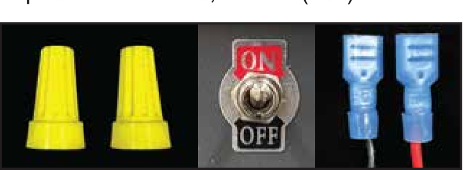 | | | **1 set** | in ammo box | connect components to power and manual switches |

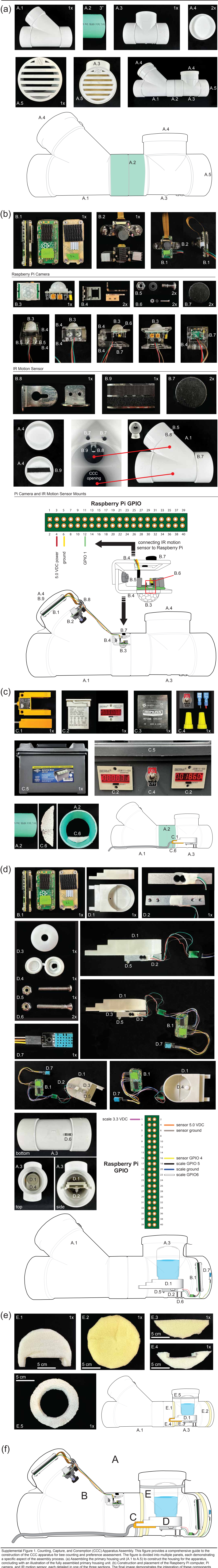


Supplemental Figure 1. Counting, Capture, and Consmption (CCC) Apparatus Assembly. This figure provides a comprehensive guide to the construction of the CCC apparatus for bee counting and preference assessment. The figure is divided into multiple panels, each demonstrating a specific aspect of the assembly process. (a) Assembling the primary housing unit (A.1 to A.5) to construct the housing for the apparatus, concluding with an illustration of the fully assembled primary housing unit. (b) Construction and placement of the Raspberry Pi computer, Pi camera, and IR motion sensor, each detailed in one of the three sections. The final image demonstrates the integration of these components within the system, including the connection between the IR motion sensor and Raspberry Pi GPIO.(c) Assembly of the IR beam counter and its protective housing unit, along with display screens. The final illustration displays the placement of these components within the system. (d)Construction of the weight scale, temperature, and humidity sensors, with images illustrating the components, their placement, and their connection to one another. The final illustration exhibits the arrangement of these components within the system and their connections to the Raspberry Pi. (e) Display of foam components used for supporting certain equipment and preventing bee entering into the rear of the apparatus. (f) A comprehensive illustration of the fully assembled CCC apparatus, with letters corresponding to the components of each panel in the preceding sections.

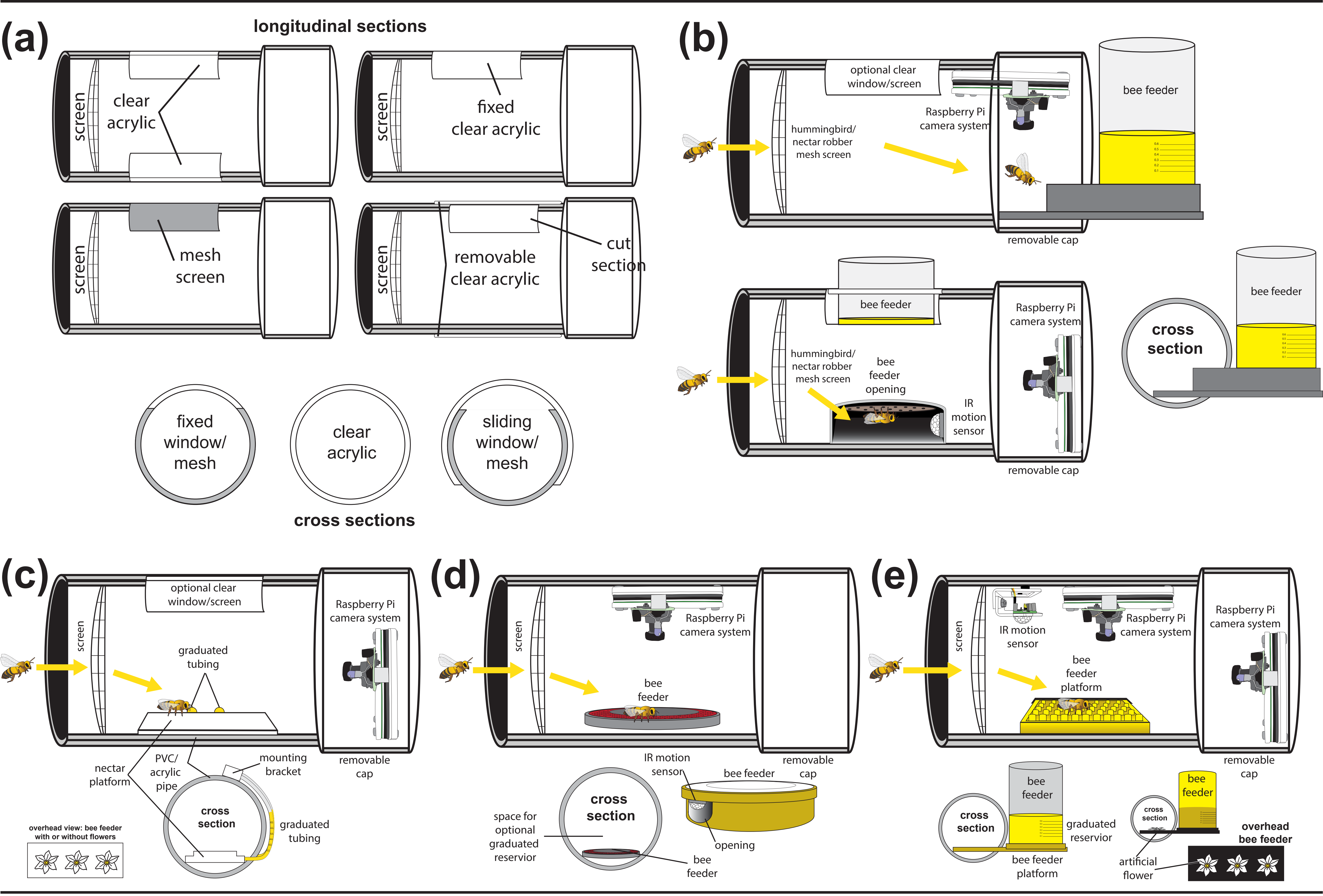


Supplemental Figure 2. Modular Alternatives to the CCC Apparatus Design. These alternative designs provide flexibility and customization options for the CCC Apparatus, catering to different research requirements and available resources. (a) showcases different housing designs for the CCC Apparatus, with options for screens to prevent unwanted visitors or acrylic for easy viewing inside the tube. (b) presents two alternative designs where the feeding apparatus is placed outside the feeding tube, with only the entrance of the feeding platform housed inside. (c) displays an alternative with a smaller volume and a small tray for bees to distinguish from, allowing volume observation from outside the apparatus. (d) illustrates two methods using standard bee feeding dishes. (e) demonstrates an alternative with the feeding solution tank outside the tube, while still having the feeding platform within.

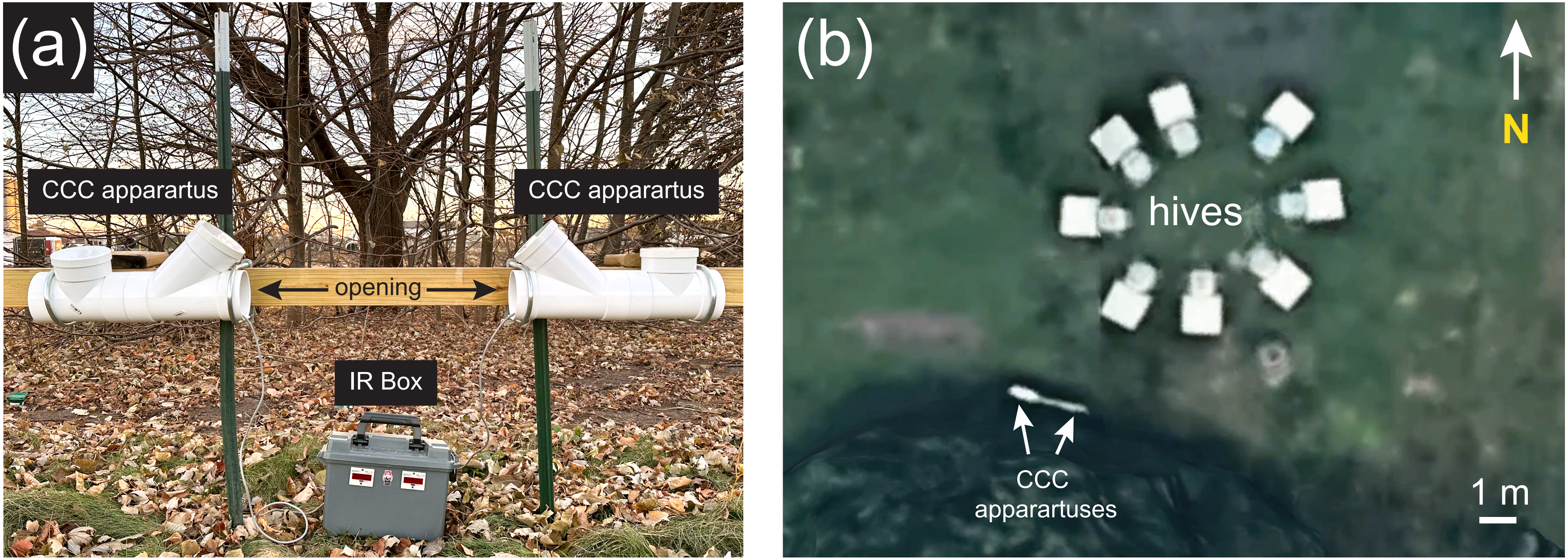


Supplemental Figure 3. Experimental Setup and Field Deployment of the CCC Apparatus (a) Photograph illustrating the field deployment of the CCC apparatus. Two apparatuses are securely mounted on a lumber board supported by T-posts, strategically positioned in proximity to the beehives. The power box and digital readouts of the Infrared (IR) beam counters are shown between the two apparatuses. (b) Google Earth image displaying the spatial arrangement of the CCC apparatuses in relation to the beehives. The geographic positioning provides an overview of the experimental site, highlighting the placement of the apparatuses for monitoring and analyzing foraging patterns among the bee colonies.

(a)

| time (min) | °C | RH |
|---|---|---|
| 0 | 12 | 88% |
| 30 | 13 | 86% |
| 60 | 14 | 84% |
| 90 | 17 | 82% |
| 120 | 17 | 74% |
| 150 | 18 | 69% |
| 180 | 20 | 64% |
| 210 | 22 | 58% |
| 240 | 22 | 52% |
| 270 | 22 | 49% |
| 300 | 22 | 49% |
| 330 | 21 | 47% |
| 360 | 21 | 46% |

(b)

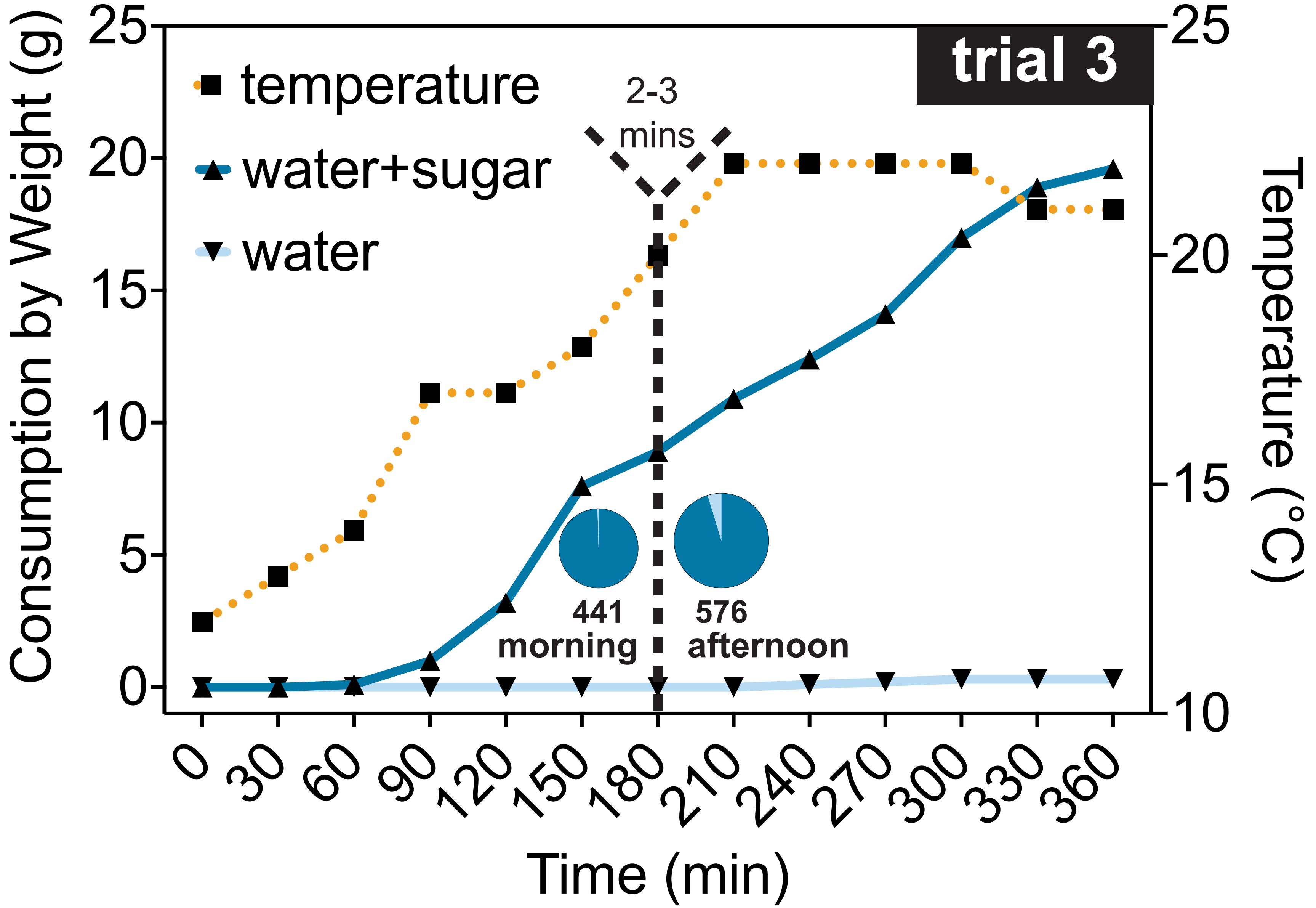


Supplemental Figure 4. Environmental Data Collection and Analysis with Consmption. (a) Temperature (°C) and relative humdidty (RH) recorded during trial 3. (b) Comparing consumption rates to ambient air temperature. The consumption rates of water+sugar (dark blue line) and water alone (light blue line) plotted along the left y-axis. The temperature (orange dotted line) is plotted on the right y-axis. The black dotted line identifies the point at which the tubes were switched from their original orientation. Comparing the consumption rates to the temperature showed that there was a significant correlation (p=<0.0001) between consumption and temperature.
